# Incorporating weather and host abundance in an iterative subseasonal-to-interannual ecological forecast system for *Ixodes scapularis*, the vector of Lyme disease

**DOI:** 10.1101/2025.10.02.677385

**Authors:** John R. Foster, Shannon LaDeau, Richard S. Ostfeld, Michael C. Dietze

## Abstract

Forecasting the population dynamics of disease vectors is critical for mitigating the risks of vector-borne diseases under a changing climate. We evaluate an iterative Bayesian forecast model of black-legged tick (*Ixodes scapularis*) phenology and population dynamics at near-term to interannual (e.g. 12 month) scales. The black-legged tick is the vector of *Borrelia burgdorferi*, the causative agent of human Lyme disease. Our forecasts consistently outperformed seasonal null models based on historical day-of-year averages, particularly during peak questing periods when disease risk is highest. Iterative data assimilation improved forecast performance over time, demonstrating the ability to adaptively learn about climate-driven shifts in demographic parameters, and reinforcing the value of long-term data to support management. Weather and climate variables emerged as the dominant predictors of nymph survival, with daily maximum temperature displacing humidity as the strongest predictor as the iterative forecast cycle evolved over time. Short-term forecasts driven by local weather observations were more accurate than those relying on seasonal climate forecasts, highlighting the importance of fine-scale weather dynamics and data for subannual predictions. At interannual scales, seasonal climate forecasts and vertebrate host (mouse) abundance were important for maintaining strong predictive skill in forecasting nymphal tick abundance, which is often used as a proxy for risk of human exposure to tick-borne disease, but forecasts were largely unaffected by larval abundance. Investment in monitoring efforts should prioritize observations of the nymphal stage to reduce forecast uncertainty. These results offer a path forward for operationalizing ecological forecasts of tick populations under environmental change and underscore the importance of adaptive, process-based models for managing tick-borne disease risk in a changing climate.

**Data availability statement:** The data that support the findings of this study are openly available. The data from the Cary Institute of Ecosystem Studies are located on Figshare. Specifically, the tick data (Ostfeld and Oggenfuss 2023) is at doi.org/10.25390/caryinstitute.23611374.v1, the mouse data (Ostfeld et al. 2024) is at doi.org/10.25390/caryinstitute.25742778.v1, and the meteorological data (Kelly, 2020) is at doi.org/10.25390/caryinstitute.11553219.v6. The NMME system data are openly available at https://www.earthsystemgrid.org/search.html?Project=NMME. Code for this manuscript (Foster, 2025) is available from Zenodo at doi.org/10.5281/zenodo.17161317.

## INTRODUCTION

Tick-borne disease (TBD) incidence is increasing across North America, with a doubling in the number of human cases of Lyme disease (LD) over the past decade serving as the most prominent example (Rosenberg et al. 2018; CDC 2021). The rise in human LD is linked to the ecology of *Ixodes scapularis*, the black-legged tick, which is the primary LD vector in eastern North America. This tick’s population dynamics emerge from a complex ecological system involving hosts, habitat, and climate (Barbour and Fish 1993; Leighton et al. 2012; Ostfeld et al. 2006; 2018). Among the many variables contributing to human risk, the density of nymphal ticks is one of the most direct and measurable proxies for LD risk (Eisen and Eisen 2024; Falco et al. 1999; Pepin et al. 2012). Because TBD risk varies dramatically in space and time (Price et al. 2024), and is continually shifting with climate and land use change (Diuk-Wasser et al. 2020; Gilbert 2021), reliable forecasts are needed to minimize human TBD risk. Risk mitigation becomes increasingly important as climate and land use change are likely to continue to drive rising TBD incidence in the coming decades (Couper et al. 2021).

Mediating TBD risk requires managing human exposure to ticks, which requires knowing when ticks are abundant in locations accessed by humans (Eisen and Eisen 2016; Stafford et al. 2017). Specific forecast models can inform risk mediation at both long and short-term horizons. Tick projections over decadal scales can be useful for assessing the geographic expansion of vectors to inform policy changes related to land use patterns (Diuk-Wasser et al. 2020; Ferraguti et al. 2023; Kopsco et al. 2023; Ogden et al. 2006). However, actionable decision-making largely occurs on shorter time scales and requires models that can generate forecasts at subseasonal to annual time horizons (e.g. 12 months in advance) and accommodate changing conditions (e.g., weather) at this same temporal scale. On the subseasonal scale, forecasts need to capture the onset, magnitude, and duration of tick activity to optimize alerts to when tick risk is highest, as the questing period (the time of year ticks are actively searching for hosts) can be measured in terms of weeks (Levi et al., 2015). On the seasonal to annual scale, forecasts of tick abundance could be used to generate tick-mitigation strategies several months in advance.

Beyond these management issues, near-term forecasts are, by their nature, verifiable as data collection is ongoing. This feature has the potential to enhance our scientific knowledge due to the iterative comparison between our system understanding (i.e. forecast) and reality (i.e. data), allowing for iterative model updates (Dietze 2017a; Lewis et al. 2023).

However, forecasting population dynamics in a changing climate is challenging, particularly when dealing with the ecological complexity of TBD systems. Weather influences development rates, survival, phenology, and questing behavior, all of which vary across tick life stages (Fouque and Reeder 2019; Gilbert et al. 2014; Levi et al. 2015; Ogden et al. 2005). For example, temperature and precipitation affect overwinter survival and molting success of nymphs (Brunner et al. 2023), low relative humidity (RH) can cause desiccation of larvae (Ginsberg et al. 2017), and soil temperature affects adult survival (Bertrand and Wilson 1996). These variables also affect the questing period, and thus the likelihood of ticks encountering humans and spreading LD (Arsnoe et al. 2019; Burgdorfer et al. 1982).

Host availability not only affects tick survival and development (Keesing et al. 2009; Ostfeld et al. 2001; 2018), but also modulates pathogen transmission cycles (Vuong et al. 2017), as certain hosts serve as competent reservoirs while others dilute infection risk (LoGiudice et al. 2003). Additionally, host populations (e.g., *Peromyscus leucopus*, the white-footed mouse) are themselves affected by climate and land use (Ostfeld & Brunner, 2015), which can create nonlinear and often unintuitive tick population responses, particularly as climate change alters the timing and intensity of these interactions (Hamer et al. 2012; Savage and Moore 2024). Finally, biotic and abiotic drivers are often nonstationary, meaning their influence on tick dynamics may vary through time or space (Rollinson et al. 2021). In the face of such complex interactions it is important to understand how the quantity and quality of information about ticks, hosts, and climate affect the predictability of tick populations. Given finite monitoring resources, such information is also critical to help optimize TBD monitoring strategies.

Generating short-term forecasts requires a modeling framework that explicitly incorporates uncertainty, particularly from weather forecasts (Deser et al., 2012), demographic stochasticity (Petchey et al., 2015), environmental stochasticity (Gauthier et al., 2016), and model structural uncertainty (Zylstra & Zipkin, 2021). Bayesian hierarchical models are well-suited to meet these needs by incorporating drivers and process dynamics (Foster et al. 2024; LaDeau et al. 2011) to generate probabilistic predictions that propagate uncertainty and iteratively integrate new data (Wikle, 2003; Dietze et al., 2018).

Previously, we introduced a process-based, stage-structured matrix model of black-legged tick abundance (Foster et al. 2024). The model integrates host abundance and daily weather data to generate daily predictions of larva, nymph, and adult abundances at a long-term research site. This model was effective at capturing daily dynamics within the calibration period, however, key advances are still needed before the model can be used to produce forecasts. Incorporating weather forecasts (and their uncertainty) of the key drivers of tick survival is a critical next step, as is the infrastructure to iteratively update both forecasts and model parameters as new data (i.e. tick drags) are collected.

This study builds on the foundation in (Foster et al. 2024) by evaluating the predictability of nymphal black-legged ticks with respect to (1) weather (and changing weather influences throughout the forecast period), (2) host abundance, and (3) larval tick abundance. Specifically, we focus on forecasts for black-legged tick populations at the Cary Institute of Ecosystem Studies (Cary), in Millbrook, NY. This location houses a long term monitoring program (Ostfeld et al. 2001; 2018) in a region where LD has been endemic for decades (CDC 2021; Ginsberg et al. 2021). We assess predictability by evaluating forecast skill across different lead times (how many days into the future the forecast is made) and by quantifying the forecast limit, which is the lead time at which the process-model becomes less skillful than the average seasonal cycling of tick abundance for each life history stage (i.e. average abundance given day-of-year). This is a strong null model as it represents the current best information that any manager could have access to without a forecast. Given that it is constructed using a particularly long, high-quality, and site-specific monitoring data set, this null is better than the information almost any manager would have in practice.

## METHODS

### Data and Site description

Three field sites at the Cary Institute of Ecosystem Studies (Green Control, Henry Control, Tea Control, hereafter referred to as Green, Henry, or Tea) were used for tick, mouse, and weather observations. Tea is about 500m from Henry, and both are >1500m from Green, centered around 41.7851N, -73.7338W. All sites are 2.25 hectares and are post-agricultural oak-dominated forest stands. Henry and Tea have been monitored since 1991, Green since 1995, with tick and mouse sampling occurring every two to four weeks during the active season from April to November each year, meaning tick sampling occurred twice during the seasonal peak in host-seeking activity by each life stage (Ostfeld et al. 2001; 2018).

### Tick and mouse data

Briefly, the stage-specific tick data (*I. scapularis*) were derived from tick drags conducted at Cary. Ticks were sampled by dragging a 1 square meter cotton cloth along the ground on three randomly chosen transects, for a total of 450 m^2^ sampled per site. Ticks were counted every 30 meters. Drags were conducted every three to four weeks between spring and late autumn. For more complete descriptions of the long-term tick monitoring at Cary see (Brunner and Ostfeld 2008; Ostfeld et al. 2001; Schauber et al. 2005). The tick data from 1995-2005 were previously used in (Foster et al. 2024) for the initial development, calibration, and validation of the tick population model. In the present study, data from 2006-2021 were used as part of our iterative forecasting design.

Mouse (*P. leucopus*) populations were monitored using a mark-recapture method, concurrent with tick monitoring (Ostfeld et al. 2001; 2018). An 11 × 11 grid of Sherman live traps was used, with stations spaced 15 m apart and two traps per station. Traps were set for two consecutive nights every three to six weeks and baited with oats. Upon first capture, mice were marked with a numbered ear tag.

Trapping did not occur during winter months. The minimum number of mice alive (MNA) for each site was directly inferred from the mark-recapture data, where if a mouse was captured on day 1 and day 3, it was inferred to be alive on day 2.

### Cary Meteorology

Meteorology was collected from Cary’s environmental monitoring program. The meteorological station is situated in a level, open field central to the three sites, at an elevation of 128 meters at N41.785823, W073.741447. Precipitation was collected with a Belfort Instrument Universal Recording Rain Gauge Series 5-780 located three meters above the ground from 1995-July 2007, and using a Geonor Precipitation Gauge Model T-200B from July 2007-2021. RH was collected with a Phys Chem Corp.

PCRC-11 or PCRC-55 (1995-1997) and a Campbell Scientific, Inc. HMP45C (1997-2021), and temperature was collected with a Campbell Scientific model 107 or 207 (1995-1998) and a Campbell Scientific, Inc. HMP45C (1998-2021). See (Kelly 2020) for a complete description of Cary’s environmental monitoring program.

### North American Multi-Model Ensemble

To investigate the feasibility of a true, year-long tick abundance forecast, we needed a seasonal weather forecast rather than the observed meteorology. We used the North American Multi-Model Ensemble (NMME) as our weather driver, as it provides a 12-month weather forecast with 10 ensemble members, and thus provides a representation of forecast uncertainty. NMME is an archived weather forecast produced monthly at a one-degree resolution and a 6-hour time step (Kirtman et al. 2014), and was used to represent the weather forecast that would have been “in-hand” at the time each tick forecast was issued. Incorporating and estimating the uncertainty across ensembles (see Supplementary information 1.1, and Figure S3.1 as an example forecast) enabled us to propagate driver uncertainty into the forecast (Dietze 2017b).

### Population Model and Iterative Updating Framework

We simulated tick population dynamics using a daily-timestep stage-structured model (Foster et al., 2024). The four stages (questing larvae, dormant nymphs, questing nymphs, and questing adults) were chosen because including dormant nymphs, a completely latent state, improved model accuracy and predictive capacity. Daily survival rates for each questing stage were modeled by four weather variables (maximum temperature, maximum RH, minimum RH, and precipitation), with weather variables standardized to their historical mean and standard deviation. Stage transitions were density-dependent on small mammals, which act as key hosts for immature ticks (Ostfeld et al. 2018). The model explicitly defines phenology by aligning empirically determined questing periods to cumulative growing degree days, with transition rates to questing states being zero during dormant periods and non-zero during questing periods (Ogden et al., 2008). Posterior parameter distributions from Foster et al. (2024) are based on a Bayesian calibration to the Cary plots using data from 1995-2005 and were used as priors in the current study.

The process-based model from (Foster et al. 2024) was used in an iterative forecasting framework (White et al. 2018; Taylor and White 2020) that ran from 2006 to 2021. This allowed us to continuously challenge the out-of-sample predictions made by the model with new data. Each new data point was first used to validate earlier predictions, and then assimilated into the model, updating both the model’s state variables (tick densities) and parameters, which then served as the basis for the next forecast. Each forecast, then, represents the current hypotheses about how this TBD system works, especially as it pertains to climate and tick survival, and each iterative prediction represents a new quantitative evaluation of our current understanding of the system.

Data processing and analysis were done in R v4.0.2 (R Core Team 2020), and hindcast simulations were carried out in NIMBLE (de Valpine et al. 2017; 2022). When using the observed weather data from Cary, the forecast horizon was 365 days. This process continued throughout the hindcasting period (2006 – 2021). However, for the years 2018-2021, we ran a parallel set of hindcasts that used the NMME weather forecasts to derive stochastic driver variables. The NMME product became available in 2018, which is why these forecasts started in 2018 instead of 2006. The hindcast involved two *concurrent* steps: a forecast step (i.e. making the prediction), and an analysis step (i.e. data assimilation).

#### Forecast step

The workflow for the hindcast is as follows: the first hindcast in 2006 was initialized 31 days before the first tick drag that year, which was on 2006-06-19 at Henry, 2006-07-03 at Green, and 2006- 07-05 at Tea. The initial condition was set using the historical mean abundance from the calibration period (1995 – 2005) for the month of the first hindcast (i.e. May at Henry and June at Green and Tea) and ran for 365 days. The next hindcast started on the day of the next tick observation (2006-08-08 at Henry and Tea, 2006-08-16 at Green), projected forward another 365 days, and used the previous hindcast’s posterior estimate of each tick-stage on the day of the observation as the initial condition (the analysis step). Each subsequent forecast repeated this pattern, being initiated on the date of the next tick observation, from the most recent posterior prediction, and ran forward another 365 days.

To increase computational efficiency, tick populations were predicted daily from the previous day, instead of across days as done in (Foster et al. 2024),

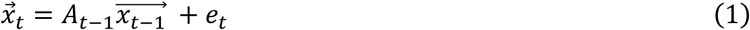

where on day *t-1,* **A** is the daily transition matrix, et is a Normally distributed process error (i.e. unexplained variability not captured by the model), and *x* is the estimated latent state vector, which was used to predict the next days’ (*t*) expected tick population *x*. For the hindcasts that used NMME, there is the added step of estimating the latent NMME variables as described in Supplementary information 1.1.

The tick hindcasts were run for each site, and model parameters were used to hindcast ticks at the calibration site (i.e. parameters calibrated at Henry were used to forecast ticks at Henry). We then ran across-site forecasts (i.e. parameters calibrated at Henry were used to forecast ticks at Green), to get an initial assessment of the model’s ability to transfer across sites. The parameter distributions for each iterative forecast were updated using the previous hindcast’s parameter distributions as priors (Supplementary information 1.2).

#### Analysis step

The analysis step was run iteratively at each time point where new tick data became available and was used to update both the state estimates of the tick population densities for each life stage and the model parameters, with these posteriors then used to initialize each subsequent forecast. At each such time point the latent tick abundance (*x*) was assumed to be normally distributed with variance *σ_x_*^2^(i.e. forecast uncertainty, a combination of initial condition, driver, parameter, and process errors), and the observed tick count data (*y*) were assumed to be Poisson distributed

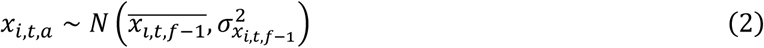

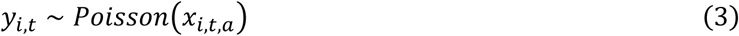

where *i* indicated life stage, *a* indicated the analysis estimate, and *f-1* indicated the previous forecast. Therefore, *x*_*i*,*t*,*a*_ becomes the first estimate of latent abundance in the new forecast, constrained by the observed tick abundance (*y*_*i*,*t*=1_) *and* informed by the previous forecast with propagated uncertainty. Parameters are updated via their influence on the previous forecast. The NIMBLE code for an individual forecast and analysis iteration is a modified version of the previous Bayesian state space calibration code (Foster et al. 2024), updated to run for a single observation time point, rather than retrospectively over a calibration time series. The MCMC for each forecast was three chains of 20K iterations, convergence was assessed using the coda package (Plummer et al. 2006), and 2K random samples with replacement were used for subsequent analysis.

### Data and sampling design experiments

We ran three sets of data inclusion experiments, which involved running parallel forecasts that included or excluded certain data.

For the NMME data experiment, we aimed to compare the accuracy of a simulated “true forecast” driven by the seasonal weather forecast that would have been available on that day to our forecast driven by the observed weather. Such an experiment is critical to assess the likely accuracy of future forecasts (via comparison to observations) and the relative increase in predictive uncertainty (e.g., interval width) associated with using an ensemble-based meteorological driver as input. Here, we ran parallel hindcasts starting in 2018 where one model used the observed weather from the Cary meteorological station (i.e. weather was treated as known during the forecast period), and compared those to a model that assimilates the NMME forecast data product.

For the larval data experiment, we aimed to assess the information contribution provided by the larval life stage to help optimize field monitoring efforts. Observing this life stage requires non-trivial effort due to their small size and large numbers, and because larval ticks are not yet infected with *B. burgdorferi*, data on larval abundance do not provide immediate information about TBD risk. Here, we ran parallel hindcasts where larval tick drag counts were either included in the analysis or excluded by setting the value of all larval observations to NA so no information was provided during data assimilation. For these experiments the structure of the model did not change.

For the mouse data experiment, we similarly aimed to inform field monitoring efforts by assessing the information contribution provided by using the abundance of an important host for immature ticks. These forecasts included or excluded the mouse population data described above. Here, if the mouse data were included, MNA was calculated from the mark-recapture data, where MNA was used to constrain transition probabilities as in (Foster et al. 2024) (i.e. mouse abundance was treated as known during the forecast period). If MNA was not included, the model structure for tick life history transition rates changed to a phenology-only model.

### Day-of-year Model

We evaluated our process-based forecast model against a simpler “day-of-year” null model, which estimates historical mean tick abundance independent of environmental covariates. This is a suitable null model due to the consistent seasonal phenology of each life stage. In the northeastern US, black-legged tick nymphs quest from late spring, peak in early summer, and become dormant by late summer (Levi et al., 2015). While this pattern is consistent, the interannual variability in questing nymph magnitude varies substantially (Ostfeld et al., 1996). Comparing our models to this null model will determine if they can accurately predict interannual variability in tick phenology and abundance.

We constructed our null forecast using a generalized additive model (GAM) smoothed over day- of-year. This model predicts the average tick abundance for any day-of-year, including days when ticks were not sampled. We used a Poisson observation model, and the smoothing was done by a thin plate regression spline, with the default smoother and number of knots (k=10) provided by the MGCV package (Wood 2017; Wood et al. 2016). The null forecast itself was generated iteratively. For every new observation, a new null forecast was generated, meaning the number of observed counts used to smooth over increased throughout the forecast period.

### Evaluating Forecast skill

Forecasts were quantitatively assessed by the continuous ranked probability score (CRPS), often used to evaluate ecological forecasts (Lewis et al. 2022; Simonis et al. 2021; Wheeler et al. 2024). The ensemble-based form of CRPS can be thought of as consisting of two components, one that calculates mean absolute error (MAE), averaged over forecast ensemble members, as a measure of accuracy and the other that calculates the mean absolute pairwise distance among ensemble members, as a measure of precision, providing an additional penalty for forecasts that are either overconfident (too small an ensemble spread relative to the magnitude of the error) or underconfident (overly pessimistic ensemble spread). Like MAE, CRPS has the same units as the response variable and the lowest (i.e. most skillful) score is zero.

To evaluate forecast skill with respect to forecast horizon, we constructed GAMs where the x- axis was lead time and the y-axis was was either forecast CRPS (a measure of absolute skill) or the difference between forecast CRPS and the CRPS at time zero (a measure of relative loss of skill over time), and the default smoother from MGCV over forecast horizon with k=5 knots. Five knots was chosen over the default 10 to reduce the “wigglyness” in the predicted target value to ease interpretation of trends with respect to forecast horizon. Additionally, for the GAMs predicting absolute CRPS, we used a log- link function to ensure CRPS predictions were zero-bound.

## RESULTS

We focus our results and evaluation of forecast skill for nymphs because it is the life stage of greatest relevance to TBD risk (Eisen and Eisen 2024; Falco et al. 1999; Pepin et al. 2012). Results for larval and adult forecasts are in the Supplementary Information 2.1 and 2.2, respectively.

### Weather impacts tick survival

We evaluated the ability of climate variables to affect daily survival, how these effects changed over the forecasting period, and what these changes mean for predictability. Daily maximum temperature became the most influential driver of nymph survival, where the median effect changed from -0.98 to - 1.53 at Green, from -1.24 to -1.58 at Henry, and from -1.00 to -1.38 at Tea (Figure 2A). This means that a one standard deviation change of daily maximum temperature decreased the odds ratio of nymph survival, and this decrease became more pronounced over the forecasting period. The magnitude of the change in the odds ratio (the difference in the odds ratio at the end of study compared to the beginning) of survival was, at the median, 2, 1.4, and 1.3 at Green, Henry, and Tea, respectively.

**Figure 1.**
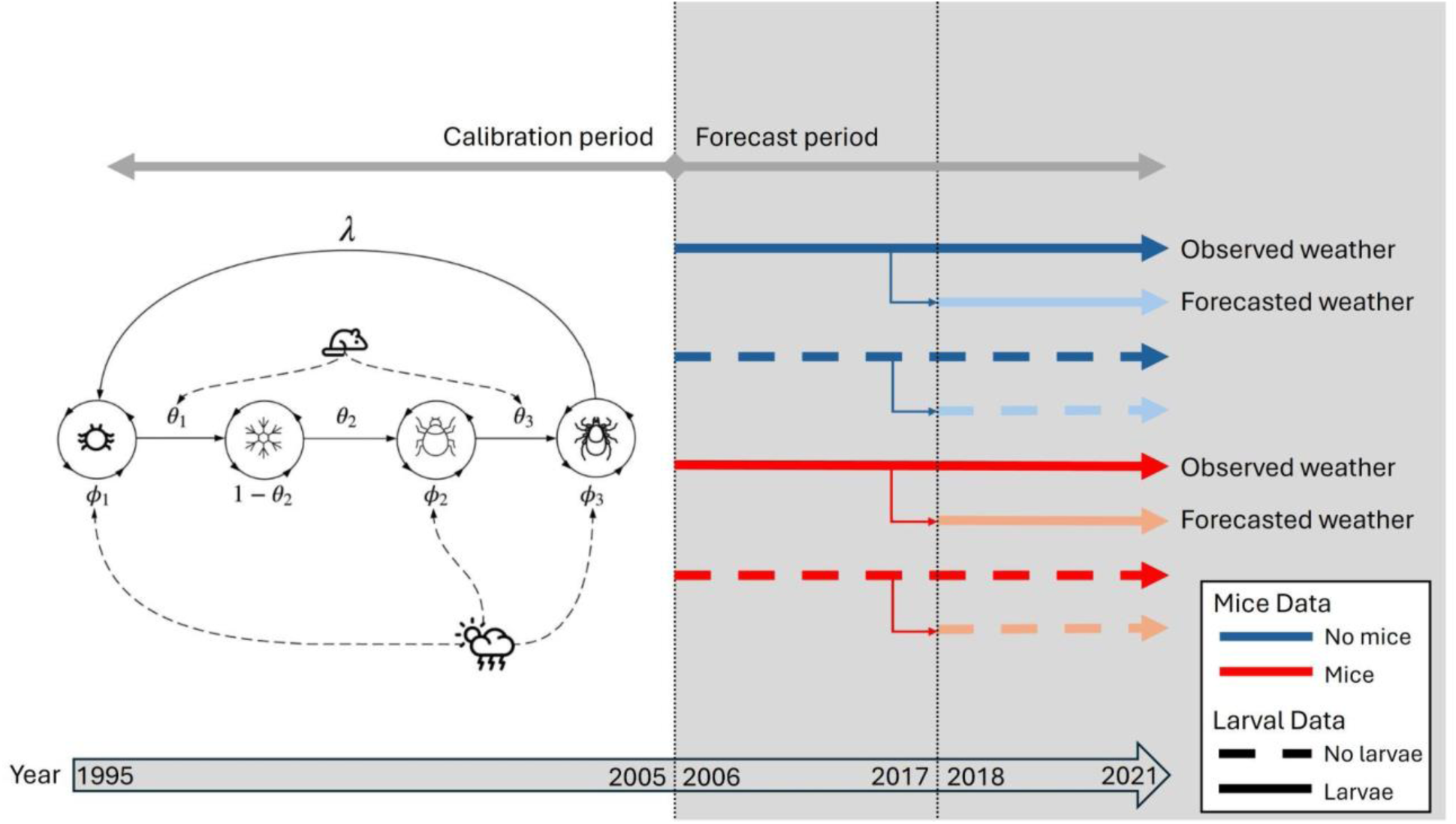
Timeline and conceptual flowchart for forecasts. The model was calibrated using data from 1995 to 2005, and is the stage structured model from Foster et al. (2024). *ϕ*_1−3_ represent the daily survival rates of larvae, nymphs, and adults, where the survival of each questing stage was constrained by daily weather. *θ*_1−3_ represent the transition rates from larvae to dormant nymph, dormant nymph to questing nymph, and questing nymph to adult where the rate from larvae to dormant nymph (*θ*_1_) and questing nymph to adult (*θ*_3_) was constrained by host abundance. *λ* is reproduction. Forecasts started in 2006 using the observed weather from the Cary meteorological station. Each solid and dashed line in the Forecast period grey area represents a forecast model fit/evaluated in this paper to evaluate the combinations of including or excluding mouse and larval field data. Starting in 2018, when the first NMME data became available, we ran parallel forecasts for each field data combination that used the forecasted weather from the NMME.

**Figure 2.**
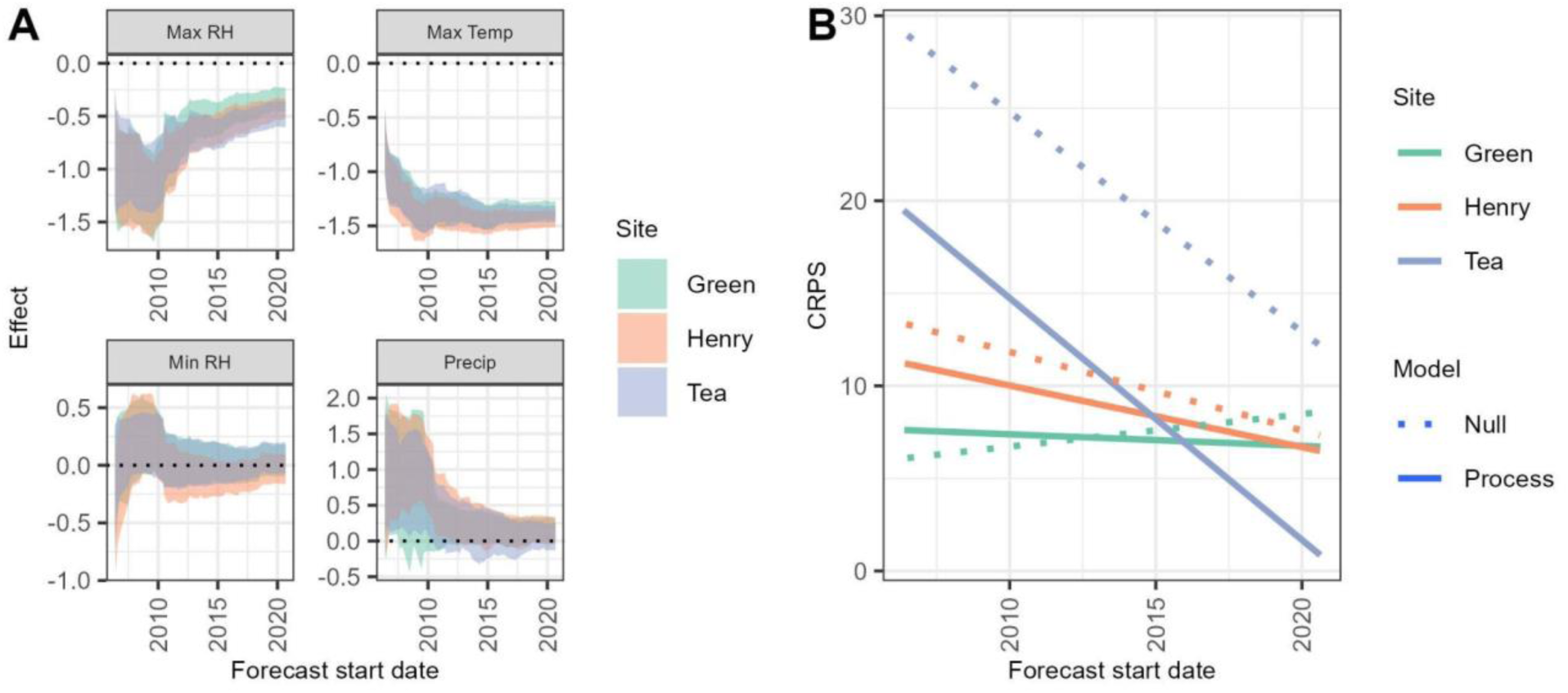
A) the change in the 95% CI of posteriors that affect nymph survival. Posteriors were updated after each forecast. B) The linear trend in forecast skill over the forecasting period at the three Cary field sites. The data shown are forecasts using the observed weather from the Cary meteorological station, larvae, and mouse data streams.

The median effect of daily maximum RH on nymph survival became less pronounced, changing from -0.83 to -0.44 at Green, from -0.95 to -0.34 at Henry, and from -0.91 to -0.41 at Tea. This means that a one standard deviation change of daily maximum RH decreased the odds ratio of nymph survival, and this decrease became less pronounced over the forecasting period. The magnitude of the change in the odds ratio of survival for daily maximum RH was -0.7, -1.2, and -1 at Green, Henry, and Tea, respectively.

The median effect of daily minimum RH on nymph survival changed minimally, from -0.04 to - 0.03 for Green, from 0.00 to 0.04 at Henry, and from -0.10 to 0.00 at Tea. The magnitude of the change in the odds ratio of survival for daily minimum RH was very small over the forecasting period, 0, 0, and -0.1 at Green, Henry, and Tea, respectively.

The median effect of daily precipitation on nymph survival became less pronounced, changing from 1.03 to 0.13 for Green, from 1.19 to 0.00 at Henry, and from 1.21 to 0.12 at Tea (Figure 2A). This means that an increase in daily precipitation increases the odds ratio of nymph survival, but this increase became less pronounced over the forecasting period, with the odds ratio declining by -1.7, -2.3, and -2.2 at Green, Henry, and Tea, respectively.

The changes in the influence of daily weather (known weather from the Cary meteorological station) on survival described above coincided with an increase in predictability (i.e. forecast skill, a lower CRPS over time) (Figure 2B). The average annual rate of change in skill for questing nymphs was 1.32 ticks/drag/year at Tea, 0.34 ticks/drag/year at Henry, and 0.07 ticks/drag/year at Green for the model that used mouse and larval data. Because the day-of-year model was likewise iteratively updated, its performance also improved at Henry and Tea by 0.4 and 1.15 ticks/drag/year, respectively, but became less skillful at Green by -0.18 ticks/drag/year. Over the course of the hindcast, this corresponds to an overall improvement (according to the linear model in Figure 2) in forecast skill by 13%, 42%, and 96% at Green, Henry, and Tea, respectively.

### Forecast skill varies seasonally

When looking at how forecast skill changed seasonally, the nymph tick forecast driven by the observed weather data consistently outperformed (lower CRPS) the day-of-year null model during the nymph questing season by up to 15 ticks/drag, but the day-of-year null outperformed the process-based forecast during the dormant season (Figure 3A). These results were insensitive to whether or not mice or larvae were included as covariates in the forecast. When driven by the NMME seasonal weather forecast, the impacts on forecast skill were modest, both during the questing and dormant periods, provided that mice data were included (Figure 3B). If mice data were excluded there was a notable drop in forecast skill during the questing season. For example, at the height of the questing season the forecasts that excluded the mouse data were on average less skillful by ∼10 nymphs/drag (Figure 3B). Similar to when the observed weather was used, there was no difference in skill comparing the forecasts that included or excluded larval data.

**Figure 3.**
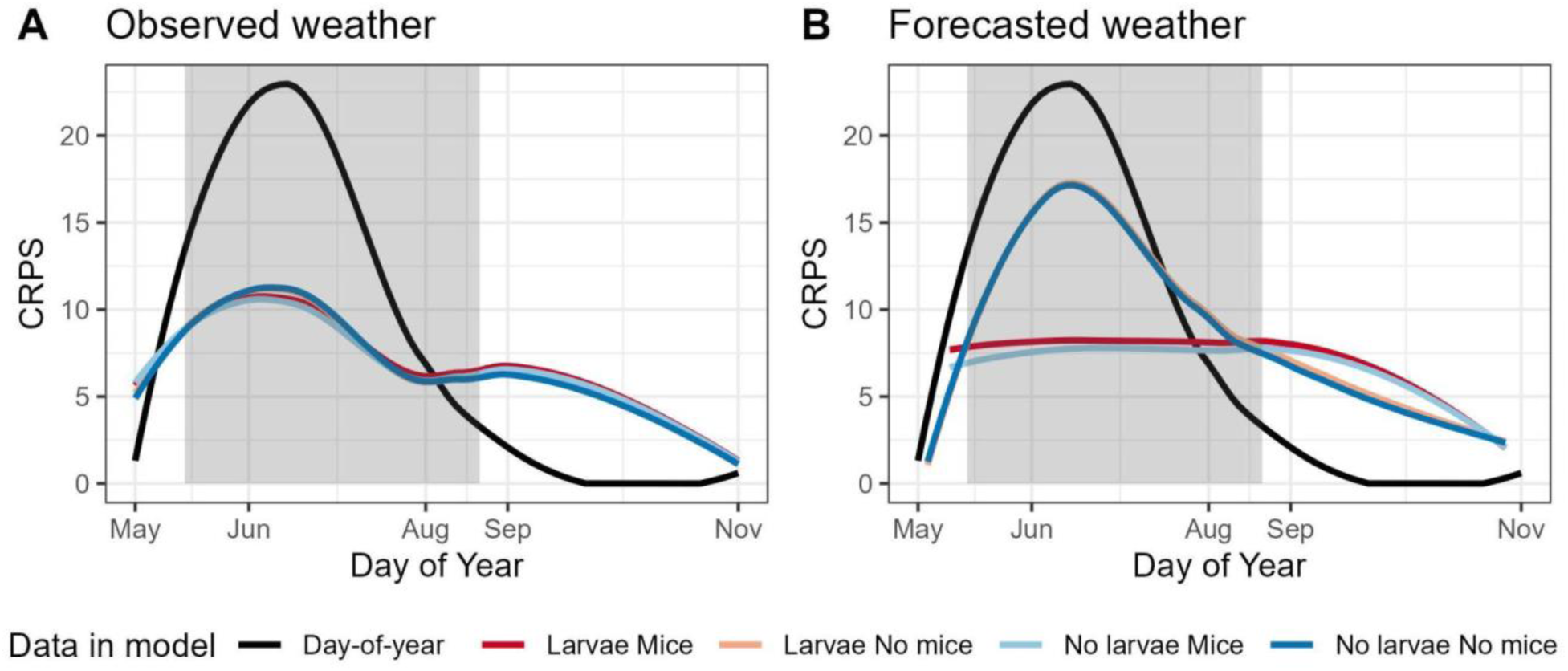
Loess curves over the continuous ranked probability score (CRPS) for nymph forecasts across all forecast horizons. A) Models that used the observed weather (Cary meteorological data) and B) models that used forecasted weather (NMME). The black line is the CRPS from the null model. The shaded region is the average start and end day-of-year for the questing period according to cumulative growing degree days over the course of the date ranges that each weather data product was used.

### Using forecasted weather reduces predictability

Here, we compare how predictability changed for tick forecasts that used the observed weather from Cary, to forecasts that used the forecasted weather from the NMME. When comparing the CRPS of the tick forecast to the null model and averaging over all of the different forecast start dates (Figure 4A and 4B) the general pattern was that the tick forecast outperformed the null model over the first month or two (i.e. subseasonal timescales), then entered a period where the null model outperformed the tick forecast, but then forecast skill subsequently returned to performing better than then null for a second time period (interannual). According to our previous definition of the forecast limit, the tick forecast thus has two forecast limits, one subannual and the other interannual.

**Figure 4.**
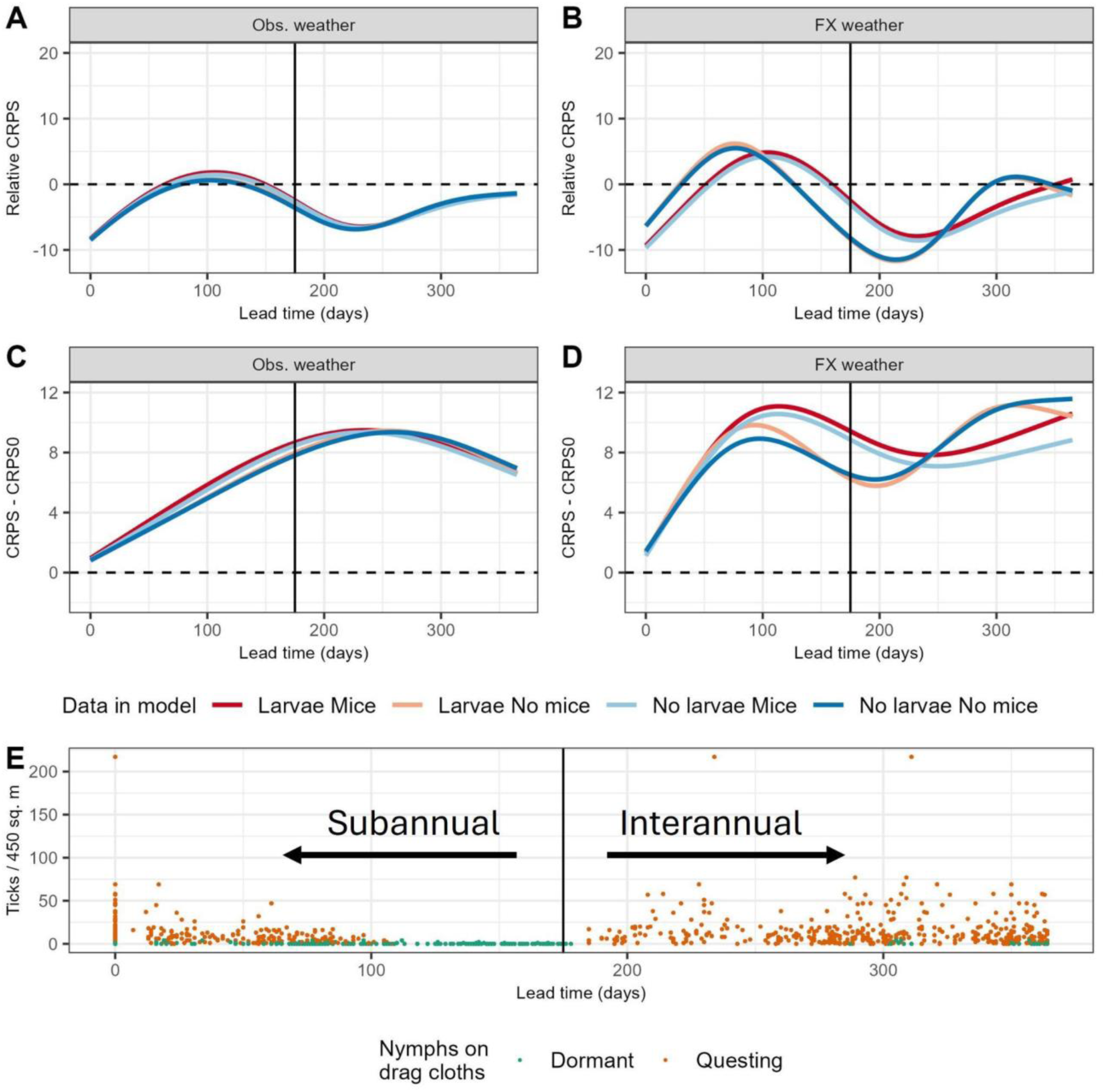
Generalized additive models (GAMs) predicting the continuous ranked probability score (CRPS) of nymph forecasts as a function of forecast lead time for questing periods only. A & B) Show GAMs predicting forecast CRPS, a measure of absolute skill, where A) depicts forecasts that used observed weather data (Obs. weather), and B) forecasts that used the forecasted weather data (FX weather). Values are relative to day-of-year forecasts, meaning values above zero indicate that the process models performed worse than the day-of-year model and values below zero indicate when the process models were more skillful. C & D) GAMs predicting the change in CRPS of nymphs relative to the “nowcast” (day 0), a measure of predictability with respect to forecast lead time, where values above zero indicate that forecasts became less predictable with lead time, below zero they got more predictable. C) Forecasts that used observed weather data, and D) forecasts that used the forecasted weather data. E) Observed nymphs on drag cloths as a function of lead time where color indicates if the tick drag occurred during the dormant (green) or questing (orange) period of the year. The vertical lines on day 175 represents the shortest lead time for interannual forecasts (the longest lead time for subannual forecasts was 178 days).

**Figure 5.**
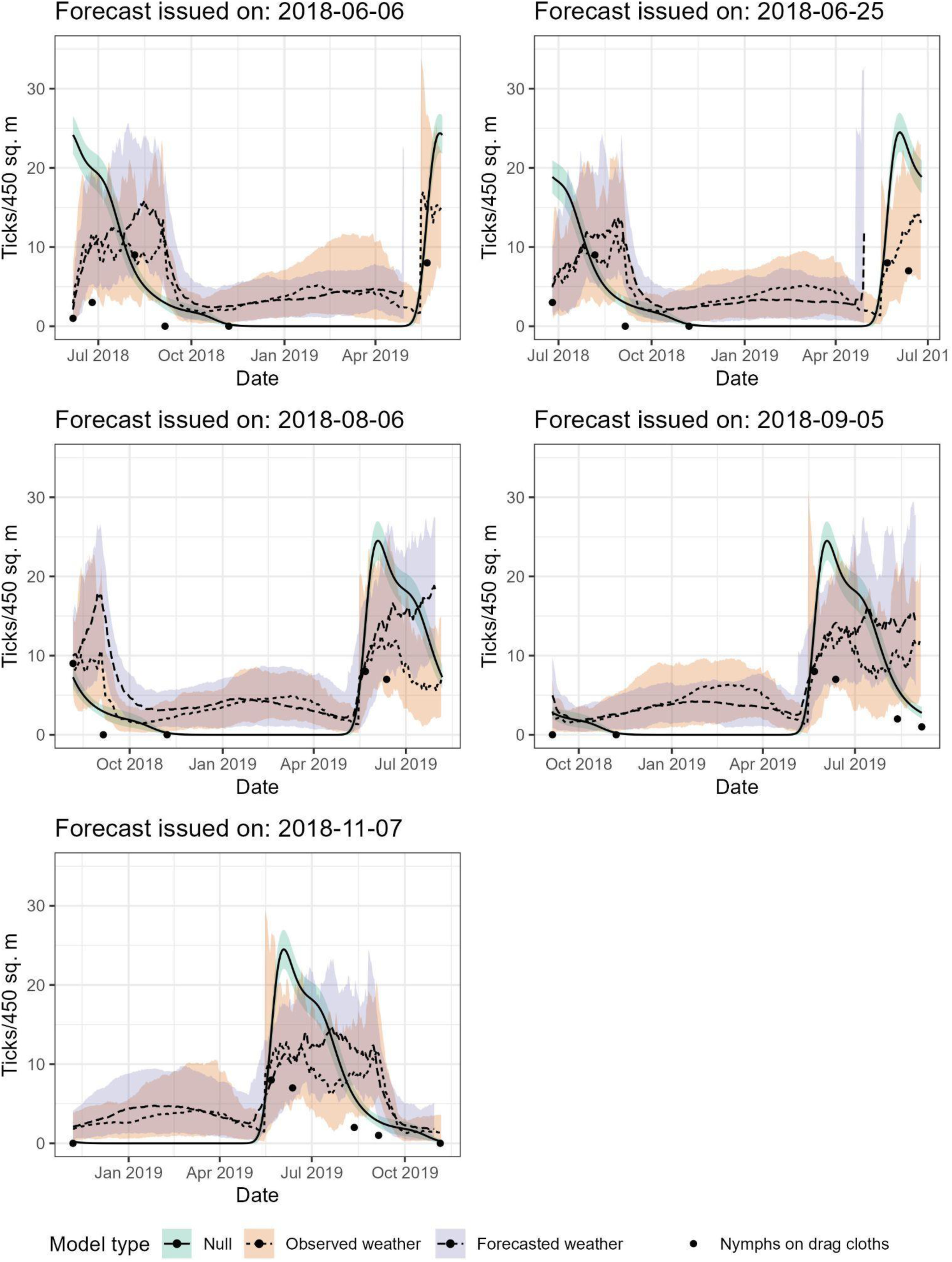
Examples of iterative forecasts issued at Green throughout 2018 for the null model (light green), the model driven by the observed weather from Cary (orange), and the model driven by the forecasted weather from NMME. The process models included mouse data and excluded larval data. Each forecast has a median (lines) and 95% predictive interval (ribbon widths). Points are nymph observations. The forecast horizon for the forecasted weather models are shorter than the observed weather models due to the horizon of the NMME data (see Supplementary information 1.1).

For subannual forecasts (i.e. forecasts that had a lead time <175 days), weather acts directly on nymphs through the instantaneous effect of weather on nymph survival. When we compared forecasts that used the observed weather data to those that used the forecasted weather data, the predictability (the number of days the process-model is no longer more skilful than the day-of-year model) for subannual forecasts declined by 13 days, from 65 days to 52 days, when mouse data was included as a covariate and 47 days, from 76 days to 29 days, when mouse data was excluded (Table 1). There was no effect of the larval data being included vs excluded with respect to the loss in predictability when comparing forecasts using the observed weather data to forecasts that use the forecast weather data.

**Table 1.**
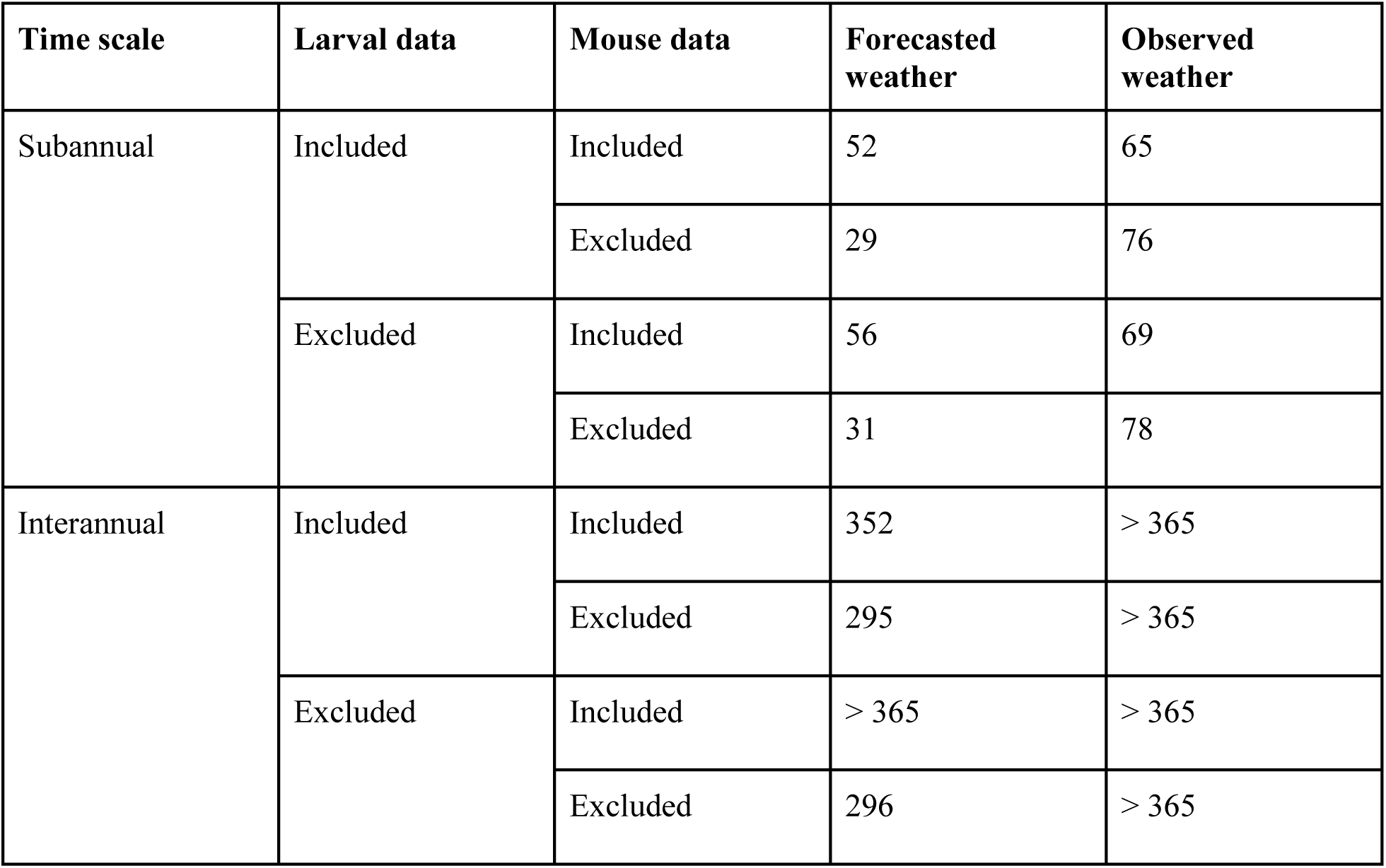
Forecast limit (lead time in days) for process-models by forecast time scale, field data included, and weather data used in the model. These are for questing nymphal ticks only. Subannual forecasts had a maximum lead time of 175 days, interannual forecasts had a minimum lead time of 178 days. The forecast limit reported is the mean forecast lead time in which the process-models became less skillful than the day-of-year model.

For interannual forecasts, weather acts directly on nymphs as above, but also integrates the compounding effect on nymph survival across seasons as well as the indirect effect on nymphs through larval tick survival and feeding success nine months earlier. For interannual forecasts, the loss in predictability for forecasts that used the observed weather data compared to those that used the forecasted weather data, when mice and larval data were included, declined from >365 days to 352 days but remained >365 days days if the mouse data were included but the larval data were excluded. The loss in predictability for interannual forecasts when the mice were excluded was at least 70 days if the larval data were included (>365 to 295) and 69 days if the larval data were excluded (>365 to 296) (Table 1).

### Including host abundance increases predictability

Host abundance (i.e. mouse data) was used in the model to affect the daily transition rate from dormant nymph (which directly follows the larval stage) to questing nymph. On the subannual scale the instantaneous effect of host abundance on nymphs is expected to be a removal mechanism by removing nymphs from the questing pool as they find hosts. Interannually, mouse abundance could act indirectly on nymphs as, over time, they increase the transition from larvae to nymph (Keesing et al. 2009).

When using the forecasted weather data, including the mouse data increased the subannual forecast limit by 23 days when using the larval data and by 25 days without the larval data. Additionally, when using the forecasted weather data, forecasts that included the mouse data were, on average, more skillful through the first 100 days of a forecast relative to forecasts that excluded the mouse data (Figure 4), corresponding to the initial questing period of nymphs during the current year. Additionally, at interannual timescales, including the mouse data increased the forecast limit by 57 days when using the larval data and by 69 days without the larval data. Forecasts including mice were also more skillful after horizon day 225 relative to forecasts that excluded the mouse data (Figure 4), corresponding to the initial questing period of nymphs during the following year.

However, when using the observed weather data, including the mouse data slightly decreased the subannual forecast limit by 11 days when using the larval data and by nine days without the larval data. Beyond that, forecasts that included the mouse data were not appreciably different in forecast skill for interannual forecast horizons relative to forecasts that excluded the mouse data (Figure 4, Table 1).

### Larval data did not impact predictability

Based on demographic theory, nymph abundance should correlate with the abundance of the preceding larval stage. Here, we determined that removing larval data had little to no effect on the median forecast skill for either questing or dormant nymphs when using the observed weather (Figure 4A), and when the forecasted weather was used the separation in forecast skill with respect to lead time was mostly due to the presence or absence of mice. Similarly, there was no appreciable difference in forecast skill as function of day-of-year or phenology (Figures 4 & 5).

For subannual forecasts using larval data and the forecasted weather, predictability, on average, fell by four days when the mice data were included and by two days when mice data were excluded, compared to forecasts that excluded the larval data. The same pattern was seen when the observed weather was used (Table 1). For interannual forecasts using larval data and the forecasted weather data, predictability, on average, fell by at least three days when mice data were included and by one day when mice data were excluded, compared to forecasts that excluded the larval data. There was no difference in predictability with respect to larval data inclusion/exclusion using the observed weather for interannual forecasts.

### Integrating the seasonal weather forecast

We were able to successfully integrate the NMME data (i.e forecasted weather) to make interannual forecasts for tick abundance (Figures 4 & 5). When comparing the full forecast model driven by forecasted weather (Figure 4B) to the equivalent forecast driven by the observed Cary meteorology (Figure 4A), we see that both show a similar pattern of decreasing skill with respect to forecast horizon. When using the forecasted weather this decline in skill (increase in CRPS) was more rapid over shorter timescales (<100 days), but the difference in skill becomes negligible beyond ∼175 days for forecasts that use mice data. Similarly, when considering seasonal changes in CRPS (Figure 3), we see that driving the tick forecast with forecasted weather does not result in an appreciable change in forecast skill in either the questing season (gray box) or the dormant season (white background). Indeed, when using the forecasted weather, CRPS is noticeably lower in the early questing season and only slightly higher in the transition from the questing season to the dormant season.

In terms of the NMME data, the forecasted weather correctly predicted the long term climatic variability for all four weather variables, where bias (forecasted weather compared to the observed weather) was centered on zero except for maximum RH, which tended to be biased low across lead times. Mean absolute error was fairly constant across lead times, with slight uptick in error after 200 days (Figure S3.2).

### Transferability

There was no appreciable difference in processed-based forecast skill for within-site (forecasts that used parameters calibrated in the same site that was forecasted) vs across-site (forecasts that used parameters originally calibrated at one site and used to forecast ticks in another) forecasts using the observed weather at subannual scales (Figure 6). Interannual forecasts using the observed weather data were, at the median, more skillful within-site than across-site for the four field data scenarios. The differences between the within-site and across-site scores were not ecologically meaningful, however, where the largest difference came from the Larvae No mice scenario where the difference was 1.41 ticks/drag (the within-site forecasts were more skillful). Forecasts that used the forecasted weather data were similarly skillful between within-site and across-site forecasts at both subannual and interannual scales. Regardless of weather data, field data, or forecast horizon, the process-based forecasts were more skillful than the day-of-year model both within and across sites (Figure 6).

**Figure 6.**
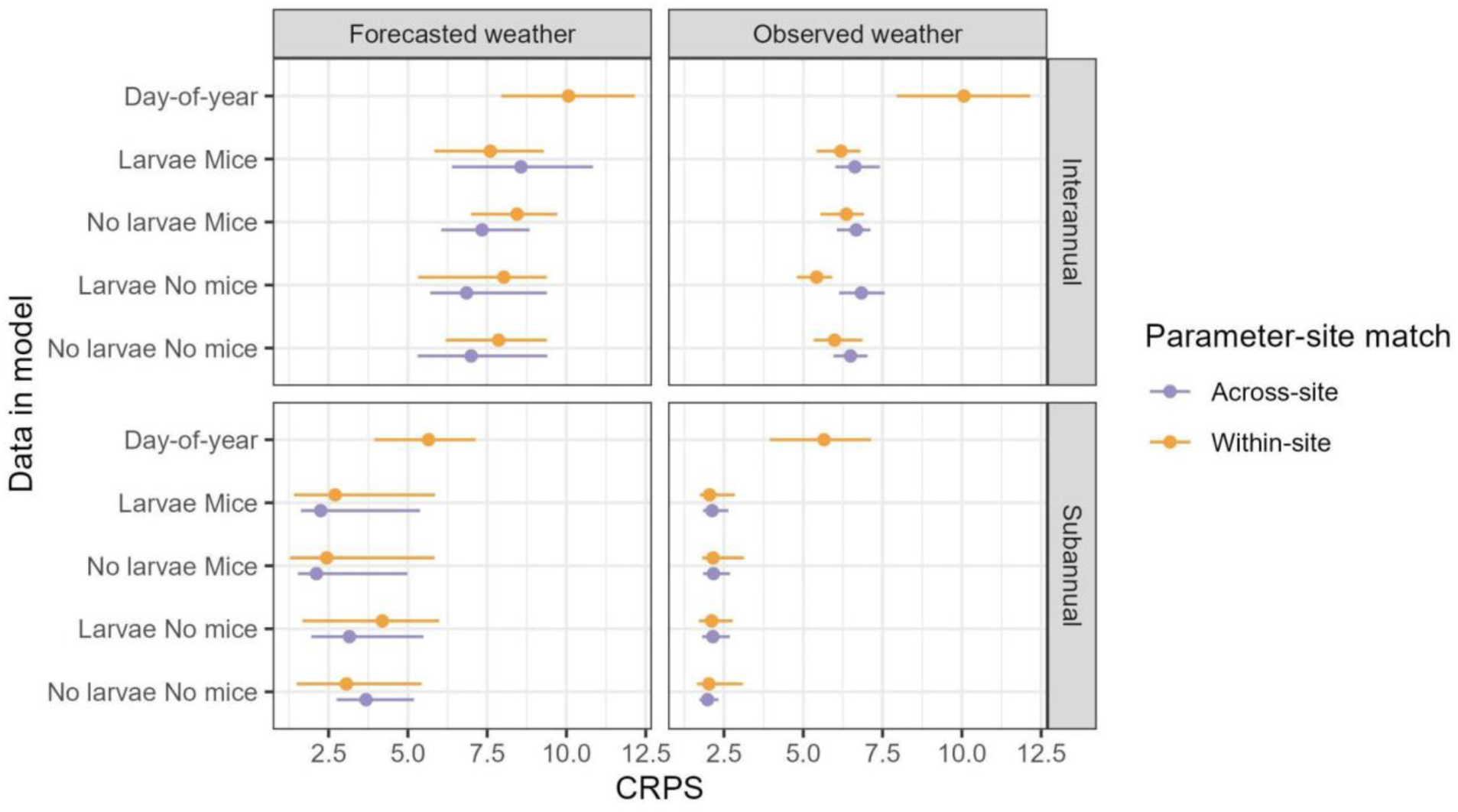
-. CRPS 90% confidence intervals and medians (x-axis) for questing nymphs given the data used in the model (y-axis), where forecasts that used parameters calibrated in the same site that was forecasted (within-site) are compared against forecasts that used parameters originally calibrated at one site and used to forecast ticks in another (across-site). The null day-of-year model was only run within-site.

## DISCUSSION

In this study we took a previously calibrated model for the population dynamics of black-legged ticks and used it to develop an iteratively-updated forecast of tick abundance and phenology. Our iterative ecological forecasts outperformed a null model that was based on long-term average tick abundance during the peak questing activity of the nymph stage, when TBD risk is highest (Arsnoe et al. 2015; Eisen and Eisen 2016; Levi et al. 2015). In other words, our process-based forecasts were more skillful than the best information currently available to managers without a forecast and are likely superior to the information most managers possess in practice.

### Weather impacts tick survival

Ticks spend >90% of their lives off host (Brunner et al. 2023; Ostfeld and Brunner 2015) and thus the impacts of weather on tick survival were expected to be a dominant factor in predicting tick population dynamics. As predicted, process-based forecasts that used the observed weather data outperformed the day-of-year forecast during the nymph questing period, successfully predicting both subseasonal and interannual variability in tick abundance and phenology. Temperature and humidity affect survival and questing behavior of black-legged ticks, and have been used to predict their abundance, development, and phenology (Eisen et al. 2016; Foster et al. 2024; Ginsberg et al. 2020; Levi et al. 2015).

The relationship between weather and tick survival changed over the course of the study (Figure 2A) allowing the model to become more skillful over time (Figure 2B). Specifically, by the end of the study, daily maximum temperature became the most influential weather variable on nymph survival, which is in contrast to our previous work which predicted that nymphs were most sensitive to daily maximum RH (Foster et al. 2024). The most likely explanation for these observations is that the final effects of weather on nymph survival reflect a more accurate and precise representation of how nymphs respond to daily and seasonal weather than at the beginning of the study, especially since the temperature and RH monitoring sensors remained unchanged throughout this period (Kelly 2020). It is also possible that how ticks respond to climatic conditions shifted over the course of the overall time series (1995- 2021). Nevertheless, ticks remain sensitive to moisture stress through the combined effects of RH and precipitation as we have previously discussed (Foster et al. 2024).

Temperature is a biologically important weather variable for nymphs, predicting their development duration (Ogden et al. 2004), and extremely high temperatures can cause desiccation (Beament 1959). Temperature has also been shown to increase time spent questing (Eisen et al. 2016; Vail and Smith 2002), which increases the chance of finding a host (Arsnoe et al. 2019; Ogden et al. 2005) therefore increasing the exposure of humans to TBDs such as LD (Eisen and Eisen 2016).

The observed changes in the effect of weather on nymph survival coincided with improved forecast skill over the course of the study, which matches theoretical expectations (Dietze 2017b) and previous ecological forecasting work (McClure et al. 2021; Niu et al. 2014; Randon et al. 2022). We extend this by showing a foreword-only MCMC data assimilation scheme can update model parameters when necessary, reducing computational burden by not refitting the entire time series with each iteration. The forecasts improved at different rates across our three sites, and we attribute this difference to the heterogeneity across sites our models have yet to explain (Oliver and Roy 2015).

### Forecast skill varies seasonally

Forecast skill showed cyclic dynamics that followed the known phenology of the nymphal black-legged tick, where the day-of-year model became more skillful at forecast lead times associated with dormant periods (Figure 3). The underlying reason for the superior performance of the day-of-year model during dormant periods likely lies in the distinction between tick abundance and tick activity. During dormant periods (molting, winter inactivity) ticks still exist in the environment, but are not measured by tick drags because they are not actively questing. The day-of-year model is trained on data that includes this seasonal cycle, but the process-based forecast of tick abundance is not capturing this distinction sufficiently and thus overpredicting dormant-season tick counts.

An immediate solution could be to combine the process-based and day-of-year forecast, switching between the two based on phenology. This suggests that the process-model may be improved via an improved representation of tick phenology and activity. Some of these issues were noted during calibration (Foster et al. 2024), but are difficult to separate from climate effects on survival when relying solely on tick drag data collected every three to four weeks, and thus suggests a need for additional data specifically around phenology and activity. Recording the time of day and microclimate variables at finer spatial scales (i.e. where the tick drag occurs) may also help disentangle activity from abundance as these factors have been shown to influence questing behavior (Dubie et al. 2018; Schulze and Jordan 2003).

It is also worth noting that this dormant season bias (abundance versus activity) is also affecting the overall relationships between lead time and skill (Figure 4). Specifically, because our forecasts are updated each time new data are assimilated, the forecast start times are based on the tick drag dates, which occur disproportionately during the active seasons. Therefore the questing period is overrepresented in subseasonal (<100 day) and interannual (>175 days) forecasts, while the dormant period is overrepresented in between, corresponding to the time period when the null model outperforms the forecast (Figure 4E). This is the reason we observed two distinct forecast limits (subseasonal versus interannual). Not only should this be kept in mind when interpreting the predictability analyses, but also suggests that explicitly accounting for dormant season inactivity will likely shift the predictability analyses to having a single forecast limit.

### Using forecasted weather reduces predictability

Forecast skill generally decreased with lead time (Figure 4C & D), which aligns with previous ecological forecasts (Randon et al. 2022; Ratnayake et al. 2019; Wheeler et al. 2024; Woelmer et al. 2022) and theory (Dietze 2017b). Furthermore, increases in CRPS with lead time tended to asymptote to a quasi-steady-state value. For forecasts driven by the observed weather, this asymptote occurred at ∼250 days and was modestly sensitive to the inclusion or exclusion of host or larval data. Forecasts driven by the NMME asymptoted to a very similar CRPS value but reached this asymptote after only ∼100 days (i.e. using the forecasted weather, instead of the observed meteorology, reduces predictability). The invariance of this skill asymptote suggests that at longer lead times the accuracy of day-to-day weather seems to be less important than seasonal trends in climate, as forecasts at this scale integrate over fluctuations in daily weather. Therefore, our forecasts suggest that the population dynamics of black-legged tick nymphs are driven by variations in climate at interannual time scales, and variations in weather at subannual scales. A possible future direction would be to explore ways to combine NMME with short-term weather forecasts, such as the 35-day NOAA Global Ensemble Forecast System, as such short-term forecast systems are likely more skillful than NMME over these timescales.

It is worth noting that forecasts for the following year (horizons >175 days) were on average more skillful than the day-of-year model, even when using the forecasted climate from NMME. Therefore, models incorporating climate forecasts and mouse populations are superior to historical averages for predicting next year’s nymph emergence and abundance. This allows land managers to more effectively target LD mitigation efforts, as nymph abundance is a better indicator of exposure risk (Gilbert, 2021).

While climatic factors remain influential in predicting questing nymph abundance, other large-scale processes which we did not consider here, such as land-use change, also affect vector abundance, confounding the relationship between climate change and vector-borne disease incidence (Franklinos et al. 2019; Medlock and Leach 2015). Accounting for these factors will be important when expanding forecasts across larger spatial scales.

### Including host abundance increases predictability

When we removed mouse data from the forecast, we found this had little effect on forecasting nymph abundance when using the observed weather, but had a large impact on both forecast skill and the forecast limit when using the NMME weather forecast (Figures 3, 4A & 5A).

Mouse abundance during questing affects questing nymph counts, because mice remove ticks from the questing pool (Ostfeld et al. 2018). We found that when the observed weather was used for subannual forecasts there was a short (11 day) decrease in predictability when mice were included (Table 1). While this decrease was surprising, given the importance of mice in regulating nymph populations in the current season, it was shorter than the mean difference between tick drags at Cary, meaning the loss in predictability is less than the number of days between observations. This result does highlight the importance of daily weather as the primary driver of nymph population dynamics in the short-term during the questing season and when finescale weather dynamics are known. When finescale weather dynamics are unavailable, or in an actual forecasting scenario when a data product like NMME must be used, host population abundance remains a major contributor to the predictability of black-legged nymph abundance with respect to forecast horizon (Figure 4) and phenology (Figure 3).

For interannual forecasts, mouse abundance in the prior larval season is expected to impact subsequent nymph abundance (about nine months later), owing to the high quality of mice as larval hosts (Ostfeld et al. 2001). Consistent with this, interannual predictability increased by at least two months when the mouse data were included and the forecasts were driven by the NMME weather forecast (Table 1). Our results suggest that when ticks begin to quest, in either the current or following year, mouse population dynamics play a crucial role in black-legged nymph population predictions under an uncertain climate.

It should be noted that the mouse abundance was treated as known during the forecast period. Therefore, these results represent a best-case scenario when using mouse abundance as we did not propagate uncertainty. At the same time, however, the no-mouse model conceptually provides a worst-case scenario, and when using NMME the no-mice model still outperformed the null model. Tick forecasts driven by mouse forecasts, e.g., based on acorn production (Ostfeld et al. 2018) are thus expected to fall somewhere in between.

### Larval data did not impact predictability

We found that removing larval data had minimal effect on the nymph predictability. This suggests that demographic forcing from larvae to nymph is minimal for the purpose of predicting the risk of encountering infection-transmitting nymphs, matching expectations (Ostfeld et al. 2006; 2018)LaDeau et al., in review, (Ostfeld et al. 2006; 2018). This conclusion could be a consequence of the extensive data collection efforts on larval abundance that were used to calibrate the model in (Foster et al. 2024), such that the larval data used in the forecasting period are simply not providing additional information, to an already well informed process, beyond that provided by nymph and adult observations. Additionally, the survival rate of larvae to the nymphal stage may be less dependent on the initial abundance of questing larvae and more influenced by weather and the availability and quality of hosts (Keesing et al. 2009; Levi et al. 2016).

If the goal is to predict nymph abundance, then the lack of information provided by the larval data suggests that tick monitoring efforts could be redirected to reduce larval sampling and instead sample nymphs more thoroughly (larger sample size = smaller observation error) or frequently. This could, in theory, improve forecast skill by reducing nymph initial condition uncertainty, which we previously demonstrated is the greatest source of uncertainty for nymph predictions during the questing period (Foster et al. 2024; Dietze 2017b).

### Future work, transferability, and operationalization

With the successful demonstration of the ability to forecast tick populations across multiple times scales, future work should move towards operationalizing these forecasts, which includes both scientific and pragmatic objectives.

Scientifically, an immediate next step is to develop a forecast for mouse abundance because in an operationalized workflow, future mouse abundance will be unknown. These could be phenomenological (Dumandan et al. 2024) or mechanistic (Wang et al. 2021). Also, to be more useful to the public, work should be done on transferring these models to other locations or to other tick species with tick drag data. This is not a small task for dynamic population models (Yates et al. 2018) because of the heterogeneity in population processes (and thus model parameters) and the labor requirements for recalibrating models.

However, there is reason for optimism here, both because of the similar parameter estimates and demonstrated transferability across sites within Cary (Figure 6), and because the iterative data assimilation algorithms already developed allow our forecast system to iteratively adapt model parameters to local conditions.

Pragmatically, implementing this forecast in an automated manner would require additional cyberinfrastructure development ((White et al. 2018). Additionally, an operational forecast system will require additional data sources, such as spatial covariates to capture habitat variability.

Disseminating forecasts to land managers and the broader public is another key area of future work. People encounter ticks not just on public lands, but through activities such as gardening or recreation. Tick control measures are often the responsibility of individuals (Piesman and Eisen 2008), meaning forecasts would need to be made publicly available. Not only will this help individuals, but hopefully it will shift responsibility from individuals to tick-management programs. Overall, development of an operational public tick forecast has the potential to mitigate human incidence of TBDs.

It should be noted, however, that we have not yet included management scenarios in our forecasts. This is a key area for future research, and would help make these forecasts more directly useful to managers. That said, any management interventions based on these forecasts that lead to changes in tick abundance would, ironically, invalidate the forecast itself. This has important implications on the continued validation and assimilation of the system unless the system had explicit management/no management counterfactuals (Dietze et al. 2018).

Finally, given that both the day-of-year null model and the process-models were able to make skillful forecasts suggests that the ecological processes represented therein are at least partially understood (Luo et al. 2011), where the day-of-year model captures average abundance and phenology and the mechanistic models link demographic processes to variations in climate and host abundance. These process-models improved over time because the parameters adapted with each iteration, and the analysis as a whole provided novel insights into the predictability of tick populations, which highlights how ecological forecasting can be used to further our understanding of natural processes (Lewis et al. 2023).

## Acknowledgements

Thanks to Cary Institute field biologists for data collection. We also thank Cary staff Deb Fargione, Cassandra Harrison, Kelly Oggenfuss, and Amy Schuler for data management, and Peter Buston, Anne Short Gianotti, Charlotte Malmborg, Tess McCabe, Kathryn Wheeler, Zoey Werbin, and Katie Zarada for manuscript feedback. Funding: National Science Foundation (DEB1947756).

## References

Arsnoe, Isis M., Graham J. Hickling, Howard S. Ginsberg, Richard McElreath, and Jean I. Tsao. 2015. “Different Populations of Blacklegged Tick Nymphs Exhibit Differences in Questing Behavior That Have Implications for Human Lyme Disease Risk.” PLOS ONE 10 (5): e0127450. 10.1371/journal.pone.0127450.

Arsnoe, Isis, Jean I. Tsao, and Graham J. Hickling. 2019. “Nymphal Ixodes Scapularis Questing Behavior Explains Geographic Variation in Lyme Borreliosis Risk in the Eastern United States.” Ticks and Tick-Borne Diseases 10 (3): 553–63. 10.1016/j.ttbdis.2019.01.001.

Barbour, Alan G., and Durland Fish. 1993. “The Biological and Social Phenomenon of Lyme Disease.” Science 260 (5114): 1610–16.

Beament, J. W. L. 1959. “The Waterproofing Mechanism of Arthropods: I. The Effect of Temperature On Cuticle Permeability in Terrestrial Insects and Ticks.” Journal of Experimental Biology 36 (2): 391–422. 10.1242/jeb.36.2.391.

Bertrand, M. R., and M. L. Wilson. 1996. “Microclimate-Dependent Survival of Unfed Adult Ixodes Scapularis (Acari:Ixodidae) in Nature: Life Cycle and Study Design Implications.” Journal of Medical Entomology 33 (4): 619–27. 10.1093/jmedent/33.4.619.

Brunner, Jesse L., Shannon L. LaDeau, Mary Killelea, Elizabeth Valentine, Megan Schierer, and Richard S. Ostfeld. 2023. “Off-Host Survival of Blacklegged Ticks in Eastern North America: A Multistage, Multiyear, Multisite Study.” Ecological Monographs 93 (3): e1572. 10.1002/ecm.1572.

Brunner, Jesse L., and Richard S. Ostfeld. 2008. “MULTIPLE CAUSES OF VARIABLE TICK BURDENS ON SMALL-MAMMAL HOSTS.” Ecology 89 (8): 2259–72. 10.1890/07-0665.1.

Burgdorfer, Willy, Alan G. Barbour, Stanley F. Hayes, Jorge L. Benach, Edgar Grunwaldt, and Jeffrey P. Davis. 1982. “Lyme Disease—a Tick-Borne Spirochetosis?” Science 216 (4552): 1317–19. 10.1126/science.7043737.

CDC. 2021. “Lyme Disease Charts and Figures: Historical Data | Lyme Disease | CDC.” May 17. https://www.cdc.gov/lyme/stats/graphs.html.

Couper, Lisa I., Andrew J. MacDonald, and Erin A. Mordecai. 2021. “Impact of Prior and Projected Climate Change on US Lyme Disease Incidence.” Global Change Biology 27 (4): 738–54. 10.1111/gcb.15435.

Dietze, Michael. 2017a. Iterative Ecological Forecasting: Needs, Opportunities, and Challenges. 94527634 Bytes. 94527634 Bytes. 10.6084/M9.FIGSHARE.4715317.

Dietze, Michael. 2017b. “Prediction in Ecology: A First-Principles Framework.” Ecological Applications 27 (7): 2048–60. 10.1002/eap.1589.

Dietze, Michael C., Andrew Fox, Lindsay M. Beck-Johnson, et al. 2018. “Iterative Near-Term Ecological Forecasting: Needs, Opportunities, and Challenges.” Proceedings of the National Academy of Sciences 115 (7): 1424–32. 10.1073/pnas.1710231115.

Diuk-Wasser, Maria A, Meredith C VanAcker, and Maria P Fernandez. 2020. “Impact of Land Use Changes and Habitat Fragmentation on the Eco-Epidemiology of Tick-Borne Diseases.” Journal of Medical Entomology, no. tjaa209 (October). 10.1093/jme/tjaa209.

Dubie, Trisha R, Justin Turner, and Bruce H Noden. 2018. “Questing Behavior and Analysis of Tick-Borne Bacteria in Ixodes Scapularis (Acari: Ixodidae) in Oklahoma.” Journal of Medical Entomology 55 (6): 1569–74. 10.1093/jme/tjy133.

Dumandan, Patricia Kaye T., Juniper L. Simonis, Glenda M. Yenni, S. K. Morgan Ernest, and Ethan P. White. 2024. “Transferability of Ecological Forecasting Models to Novel Biotic Conditions in a Long-Term Experimental Study.” Ecology 105 (11): e4406. 10.1002/ecy.4406.

Eisen, Lars, and Rebecca J. Eisen. 2016. “Critical Evaluation of the Linkage Between Tick-Based Risk Measures and the Occurrence of Lyme Disease Cases.” Journal of Medical Entomology 53 (5): 1050–62. 10.1093/jme/tjw092.

Eisen, Rebecca J., and Lars Eisen. 2024. “Evaluation of the Association between Climate Warming and the Spread and Proliferation of *Ixodes Scapularis* in Northern States in the Eastern United States.” Ticks and Tick-Borne Diseases 15 (1): 102286. 10.1016/j.ttbdis.2023.102286.

Eisen, Rebecca J., Lars Eisen, Nicholas H. Ogden, and Charles B. Beard. 2016. “Linkages of Weather and Climate With *Ixodes Scapularis* and *Ixodes Pacificus* (Acari: Ixodidae), Enzootic Transmission of *Borrelia Burgdorferi* , and Lyme Disease in North America.” Journal of Medical Entomology 53 (2): 250–61. 10.1093/jme/tjv199.

Falco, R. C., D. F. McKenna, T. J. Daniels, et al. 1999. “Temporal Relation between Ixodes Scapularis Abundance and Risk for Lyme Disease Associated with Erythema Migrans.” American Journal of Epidemiology 149 (8): 771–76. 10.1093/oxfordjournals.aje.a009886.

Ferraguti, Martina, Sergio Magallanes, Marcela Suarez-Rubio, Paul J. J. Bates, Alfonso Marzal, and Swen C. Renner. 2023. “Does Land-Use and Land Cover Affect Vector-Borne Diseases? A Systematic Review and Meta-Analysis.” Landscape Ecology 38 (10): 2433–51. 10.1007/s10980-023-01746-3.

Foster, John R., Shannon L. LaDeau, Kelly Oggenfuss, Richard S. Ostfeld, and Michael C. Dietze. 2024. “A Modified Matrix Model Captures the Population Dynamics for the Primary Vector of Lyme Disease in North America.” Ecosphere 15 (10): e70022. 10.1002/ecs2.70022.

Fouque, Florence, and John C. Reeder. 2019. “Impact of Past and On-Going Changes on Climate and Weather on Vector-Borne Diseases Transmission: A Look at the Evidence.” Infectious Diseases of Poverty 08 (03): 1–9. 10.1186/s40249-019-0565-1.

Franklinos, Lydia H. V., Kate E. Jones, David W. Redding, and Ibrahim Abubakar. 2019. “The Effect of Global Change on Mosquito-Borne Disease.” The Lancet Infectious Diseases 19 (9): e302–12. 10.1016/S1473-3099(19)30161-6.

Gilbert, Lucy. 2021. “The Impacts of Climate Change on Ticks and Tick-Borne Disease Risk.” Annual Review of Entomology 66 (1): 373–88. 10.1146/annurev-ento-052720-094533.

Gilbert, Lucy, Jennifer Aungier, and Joseph L. Tomkins. 2014. “Climate of Origin Affects Tick (Ixodes Ricinus) Host-Seeking Behavior in Response to Temperature: Implications for Resilience to Climate Change?” Ecology and Evolution 4 (7): 1186–98. 10.1002/ece3.1014.

Ginsberg, Howard S., Marisa Albert, Lixis Acevedo, et al. 2017. “Environmental Factors Affecting Survival of Immature Ixodes Scapularis and Implications for Geographical Distribution of Lyme Disease: The Climate/Behavior Hypothesis.” PLOS ONE 12 (1): e0168723. 10.1371/journal.pone.0168723.

Ginsberg, Howard S., Graham J. Hickling, Russell L. Burke, et al. 2021. “Why Lyme Disease Is Common in the Northern US, but Rare in the South: The Roles of Host Choice, Host-Seeking Behavior, and Tick Density.” PLOS Biology 19 (1): e3001066. 10.1371/journal.pbio.3001066.

Ginsberg, Howard S., Eric L. Rulison, Jasmine L. Miller, et al. 2020. “Local Abundance of Ixodes Scapularis in Forests: Effects of Environmental Moisture, Vegetation Characteristics, and Host Abundance.” Ticks and Tick-Borne Diseases 11 (1): 101271. 10.1016/j.ttbdis.2019.101271.

Hamer, Sarah A., Graham J. Hickling, Jennifer L. Sidge, Edward D. Walker, and Jean I. Tsao. 2012. “Synchronous Phenology of Juvenile *Ixodes Scapularis*, Vertebrate Host Relationships, and Associated Patterns of *Borrelia Burgdorferi* Ribotypes in the Midwestern United States.” Ticks and Tick-Borne Diseases 3 (2): 65–74. 10.1016/j.ttbdis.2011.11.004.

Keesing, F., J. Brunner, S. Duerr, et al. 2009. “Hosts as Ecological Traps for the Vector of Lyme Disease.” Proceedings of the Royal Society B: Biological Sciences 276 (1675): 3911–19. 10.1098/rspb.2009.1159.

Kelly, Vicky. 2020. “Cary Environmental Monitoring Program Daily Meteorological and Solar Radiation Data: 1988-2021.” Cary Institute, January 8. 10.25390/caryinstitute.11553219.v3.

Kirtman, Ben P., Dughong Min, Johnna M. Infanti, et al. 2014. “The North American Multimodel Ensemble: Phase-1 Seasonal-to-Interannual Prediction; Phase-2 toward Developing Intraseasonal Prediction.” Bulletin of the American Meteorological Society. Bulletin of the American Meteorological Society 95 (4): 585–601. 10.1175/BAMS-D-12-00050.1.

Kopsco, Heather L., Peg Gronemeyer, Nohra Mateus-Pinilla, and Rebecca L. Smith. 2023. “Current and Future Habitat Suitability Models for Four Ticks of Medical Concern in Illinois, USA.” Insects 14 (3): 3. 10.3390/insects14030213.

LaDeau, Shannon L., Gregory E. Glass, N. Thompson Hobbs, Andrew Latimer, and Richard S. Ostfeld. 2011. “Data–Model Fusion to Better Understand Emerging Pathogens and Improve Infectious Disease Forecasting.” Ecological Applications 21 (5): 1443–60. 10.1890/09-1409.1.

Leighton, Patrick A., Jules K. Koffi, Yann Pelcat, L. Robbin Lindsay, and Nicholas H. Ogden. 2012. “Predicting the Speed of Tick Invasion: An Empirical Model of Range Expansion for the Lyme Disease Vector *Ixodes Scapularis* in Canada: *Predicting* I. Scapularis *Invasion*.” Journal of Applied Ecology 49 (2): 457–64. 10.1111/j.1365-2664.2012.02112.x.

Levi, Taal, Felicia Keesing, Robert D. Holt, Michael Barfield, and Richard S. Ostfeld. 2016. “Quantifying Dilution and Amplification in a Community of Hosts for Tick-Borne Pathogens.” Ecological Applications 26 (2): 484–98. 10.1890/15-0122.

Levi, Taal, Felicia Keesing, Kelly Oggenfuss, and Richard S. Ostfeld. 2015. “Accelerated Phenology of Blacklegged Ticks under Climate Warming.” Philosophical Transactions of the Royal Society B: Biological Sciences 370 (1665): 20130556. 10.1098/rstb.2013.0556.

Lewis, Abigail S. L., Christine R. Rollinson, Andrew J. Allyn, et al. 2023. “The Power of Forecasts to Advance Ecological Theory.” Methods in Ecology and Evolution 14 (3): 746–56. 10.1111/2041-210X.13955.

Lewis, Abigail S. L., Whitney M. Woelmer, Heather L. Wander, et al. 2022. “Increased Adoption of Best Practices in Ecological Forecasting Enables Comparisons of Forecastability.” Ecological Applications 32 (2): e2500. 10.1002/eap.2500.

LoGiudice, Kathleen, Richard S. Ostfeld, Kenneth A. Schmidt, and Felicia Keesing. 2003. “The Ecology of Infectious Disease: Effects of Host Diversity and Community Composition on Lyme Disease Risk.” Proceedings of the National Academy of Sciences of the United States of America 100 (2): 567–71.

Luo, Yiqi, Kiona Ogle, Colin Tucker, et al. 2011. “Ecological Forecasting and Data Assimilation in a Data-Rich Era.” Ecological Applications 21 (5): 1429–42. 10.1890/09-1275.1.

McClure, Ryan P., R. Quinn Thomas, Mary E. Lofton, Whitney M. Woelmer, and Cayelan C. Carey. 2021. “Iterative Forecasting Improves Near-Term Predictions of Methane Ebullition Rates.” Frontiers in Environmental Science 9 (December). 10.3389/fenvs.2021.756603.

Medlock, Jolyon M., and Steve A. Leach. 2015. “Effect of Climate Change on Vector-Borne Disease Risk in the UK.” The Lancet Infectious Diseases 15 (6): 721–30. 10.1016/S1473-3099(15)70091-5.

Niu, Shuli, Yiqi Luo, Michael C. Dietze, et al. 2014. “The Role of Data Assimilation in Predictive Ecology.” Ecosphere 5 (5): art65. 10.1890/ES13-00273.1.

Ogden, N. H., L. R. Lindsay, G. Beauchamp, et al. 2004. “Investigation of Relationships Between Temperature and Developmental Rates of Tick Ixodes Scapularis (Acari: Ixodidae) in the Laboratory and Field.” Journal of Medical Entomology 41 (4): 622–33. 10.1603/0022-2585-41.4.622.

Ogden, N.H., M. Bigras-Poulin, C.J. O’Callaghan, et al. 2005. “A Dynamic Population Model to Investigate Effects of Climate on Geographic Range and Seasonality of the Tick Ixodes Scapularis.” International Journal for Parasitology 35 (4): 375–89. 10.1016/j.ijpara.2004.12.013.

Ogden, N.H., A. Maarouf, I.K. Barker, et al. 2006. “Climate Change and the Potential for Range Expansion of the Lyme Disease Vector Ixodes Scapularis in Canada.” International Journal for Parasitology 36 (1): 63–70. 10.1016/j.ijpara.2005.08.016.

Oliver, Tom H., and David B. Roy. 2015. “The Pitfalls of Ecological Forecasting.” Biological Journal of the Linnean Society 115 (3): 767–78. 10.1111/bij.12579.

Ostfeld, Richard, and Kelly Oggenfuss. 2023. “Long Term Monitoring of the Dynamics of Rodents, Ticks and Lyme Disease Risk in Oak Forests: Mouse and Tick Data, 1995-2005.” 10.25390/caryinstitute.23611374.v1.

Ostfeld, Richard S., and Jesse L. Brunner. 2015. “Climate Change and *Ixodes* Tick-Borne Diseases of Humans.” Philosophical Transactions of the Royal Society B: Biological Sciences 370 (1665): 20140051. 10.1098/rstb.2014.0051.

Ostfeld, Richard S, Charles D Canham, Kelly Oggenfuss, Raymond J Winchcombe, and Felicia Keesing. 2006. “Climate, Deer, Rodents, and Acorns as Determinants of Variation in Lyme-Disease Risk.” PLoS Biology 4 (6): e145. 10.1371/journal.pbio.0040145.

Ostfeld, Richard S., Taal Levi, Felicia Keesing, Kelly Oggenfuss, and Charles D. Canham. 2018. “Tick- Borne Disease Risk in a Forest Food Web.” Ecology 99 (7): 1562–73. 10.1002/ecy.2386.

Ostfeld, Richard S., Eric M. Schauber, Charles D. Canham, Felicia Keesing, Clive G. Jones, and Jerry O. Wolff. 2001. “Effects of Acorn Production and Mouse Abundance on Abundance and *Borrelia Burgdorferi* Infection Prevalence of Nymphal *Ixodes Scapularis* Ticks.” Vector-Borne and Zoonotic Diseases 1 (1): 55–63. 10.1089/153036601750137688.

Pepin, Kim M., Rebecca J. Eisen, Paul S. Mead, et al. 2012. “Geographic Variation in the Relationship between Human Lyme Disease Incidence and Density of Infected Host-Seeking Ixodes Scapularis Nymphs in the Eastern United States.” The American Journal of Tropical Medicine and Hygiene 86 (6): 1062–71. 10.4269/ajtmh.2012.11-0630.

Piesman, Joseph, and Lars Eisen. 2008. “Prevention of Tick-Borne Diseases.” Annual Review of Entomology 53 (1): 323–43. 10.1146/annurev.ento.53.103106.093429.

Plummer, Martyn, Nicky Best, Kate Cowles, and Karen Vines. 2006. “CODA: Convergence Diagnosis and Output Analysis for MCMC.” R News 6 (1): 7–11.

Price, Lucas E., Jonathan M. Winter, Jamie L. Cantoni, et al. 2024. “Spatial and Temporal Distribution of Ixodes Scapularis and Tick-Borne Pathogens across the Northeastern United States.” Parasites & Vectors 17 (1): 481. 10.1186/s13071-024-06518-9.

R Core Team. 2020. R: A Language and Environment for Statistical Computing. R Foundation for Statistical Computing. https://www.R-project.org/.

Randon, Marine, Michael Dowd, and Ruth Joy. 2022. “A Real-Time Data Assimilative Forecasting System for Animal Tracking.” Ecology 103 (8): e3718. 10.1002/ecy.3718.

Ratnayake, H. U., M. R. Kearney, P. Govekar, D. Karoly, and J. A. Welbergen. 2019. “Forecasting Wildlife Die-Offs from Extreme Heat Events.” Animal Conservation 22 (4): 386–95. 10.1111/acv.12476.

Rollinson, Christine R, Andrew O Finley, M Ross Alexander, et al. 2021. “Working across Space and Time: Nonstationarity in Ecological Research and Application.” Frontiers in Ecology and the Environment 19 (1): 66–72. 10.1002/fee.2298.

Rosenberg, Ronald, Nicole P. Lindsey, Marc Fischer, et al. 2018. “*Vital Signs* : Trends in Reported Vectorborne Disease Cases — United States and Territories, 2004–2016.” MMWR. Morbidity and Mortality Weekly Report 67 (17): 496–501. 10.15585/mmwr.mm6717e1.

Savage, Joseph D. T., and Christopher M. Moore. 2024. “How Do Host Population Dynamics Impact Lyme Disease Risk Dynamics in Theoretical Models?” PLOS ONE 19 (5): e0302874. 10.1371/journal.pone.0302874.

Schauber, Eric M., Richard S. Ostfeld, and Andrew S. Evans, Jr. 2005. “WHAT IS THE BEST PREDICTOR OF ANNUAL LYME DISEASE INCIDENCE: WEATHER, MICE, OR ACORNS?” Ecological Applications 15 (2): 575–86. 10.1890/03-5370.

Schulze, Terry L., and Robert A. Jordan. 2003. “Meteorologically Mediated Diurnal Questing of Ixodes Scapularis and Amblyomma Americanum (Acari: Ixodidae) Nymphs.” Journal of Medical Entomology 40 (4): 395–402. 10.1603/0022-2585-40.4.395.

Simonis, Juniper L., Ethan P. White, and S. K. Morgan Ernest. 2021. “Evaluating Probabilistic Ecological Forecasts.” Ecology 102 (8): e03431. 10.1002/ecy.3431.

Stafford, Kirby C, III, Scott C Williams, and Goudarz Molaei. 2017. “Integrated Pest Management in Controlling Ticks and Tick-Associated Diseases.” Journal of Integrated Pest Management 8 (1): 28. 10.1093/jipm/pmx018.

Taylor, Shawn D., and Ethan P. White. 2020. Influence of Climate Forecasts, Data Assimilation, and Uncertainty Propagation on the Performance of near-Term Phenology Forecasts. Preprint. Ecology. 10.1101/2020.08.18.256057.

Vail, Stephen G., and Gary Smith. 2002. “Vertical Movement and Posture of Blacklegged Tick (Acari: Ixodidae) Nymphs as a Function of Temperature and Relative Humidity in Laboratory Experiments.” Journal of Medical Entomology 39 (6): 842–46. 10.1603/0022-2585-39.6.842.

Valpine, Perry de, Christopher Paciorek, Daniel Turek, et al. 2022. NIMBLE: MCMC, Particle Filtering, and Programmable Hierarchical Modeling. Version 0.12.2. 10.5281/zenodo.1211190.

Valpine, Perry de, Daniel Turek, Christopher J. Paciorek, Clifford Anderson-Bergman, Duncan Temple Lang, and Rastislav Bodik. 2017. “Programming With Models: Writing Statistical Algorithms for General Model Structures With NIMBLE.” Journal of Computational and Graphical Statistics 26 (2): 403–13. 10.1080/10618600.2016.1172487.

Vuong, Holly B., Grace S. Chiu, Peter E. Smouse, et al. 2017. “Influences of Host Community Characteristics on Borrelia Burgdorferi Infection Prevalence in Blacklegged Ticks.” PLOS ONE 12 (1): e0167810. 10.1371/journal.pone.0167810.

Wang, Deng, Dean P. Anderson, Ke Li, Yongwang Guo, Zaixue Yang, and Roger P. Pech. 2021. “Predicted Population Dynamics of an Indigenous Rodent, Ap*Odemus Agrarius*, in an Agricultural System.” Crop Protection 147 (September): 105683. 10.1016/j.cropro.2021.105683.

Wheeler, Kathryn I., Michael C. Dietze, David LeBauer, et al. 2024. “Predicting Spring Phenology in Deciduous Broadleaf Forests: NEON Phenology Forecasting Community Challenge.” Agricultural and Forest Meteorology 345 (February): 109810. 10.1016/j.agrformet.2023.109810.

White, Ethan P., Glenda M. Yenni, Shawn D. Taylor, et al. 2018. *Developing an Automated Iterative Near-Term Forecasting System for an Ecological Study*. Preprint. Ecology. 10.1101/268623.

Woelmer, Whitney M., R. Quinn Thomas, Mary E. Lofton, Ryan P. McClure, Heather L. Wander, and Cayelan C. Carey. 2022. “Near-Term Phytoplankton Forecasts Reveal the Effects of Model Time Step and Forecast Horizon on Predictability.” Ecological Applications 32 (7): e2642. 10.1002/eap.2642.

Wood, S. N. 2017. Generalized Additive Models: An Introduction with R. 2nd ed. Chapman and Hall/CRC.

Wood, S. N., N., Pya, and B. S"afken. 2016. “Smoothing Parameter and Model Selection for General Smooth Models (with Discussion).” Journal of the American Statistical Association 111: 1548– 75.

Yates, Katherine L., Phil J. Bouchet, M. Julian Caley, et al. 2018. “Outstanding Challenges in the Transferability of Ecological Models.” Trends in Ecology & Evolution 33 (10): 790–802. 10.1016/j.tree.2018.08.001.

