## Supplement for "Incorporating weather and host abundance in an iterative subseasonal-to-interannual ecological forecast system for *Ixodes scapularis*, the vector of Lyme disease"

### 1 Supplementary Methods

#### 1.1 North American Multi-Model Ensemble

The North American Multi-Model Ensemble (NMME) is a seasonal weather forecast product with a 365-day horizon across ten ensemble members. For each ensemble member on each day, we calculated the daily maximum temperature, maximum and minimum relative humidity, and total precipitation. Before data assimilation, each NMME variable was standardized to the historical mean and standard deviation as described above. We then applied a bias correction to the historical mean for all four weather variables. For temperature and minimum and maximum relative humidity, we calculated the annual mean for each ensemble of each variable, then calculated the difference between this and the calibration period observed means for each variable. This bias was added to each day of each variable of each ensemble. An example of the bias correction is in supplementary Figure S3.1.

The precipitation forecasts needed a different adjustment, as most days were predicted to have greater than zero precipitation, forecasting a constant drizzle throughout the year. To correct for this, we found the average annual cumulative precipitation at Cary over the calibration period (1134.16 mm). For each ensemble and forecast, we calculated the cumulative amount of precipitation within the forecast horizon. From this cumulative value we subtracted the 1134.16 mm to get the yearly bias between the historical mean annual precipitation and the annual NMME forecast precipitation, then calculated the average daily bias from this difference. Across each forecast year and ensemble, the corrected precipitation data had a median of 140.5 days (90% CI: 82.1-176) with zero precipitation compared to 10 days (90% CI: 5-16) for the uncorrected precipitation data. For comparison, the Cary weather station recorded a median of 243 days (90% CI: 219-260) each year without precipitation.

For each day of the tick forecast, we used an error-in-variables approach to estimate the true value for each weather variable and their associated uncertainties for each day. We used a multivariate censorship model to account for the theoretical bounds of relative humidity and precipitation, which are bound between 0 and 1 and bound at zero, respectively. Temperature was not censored.

$$\mu_t \sim MVN(\mu_0, \Sigma) \quad [1]$$

$$\Omega_t \sim Wishart(\Lambda,\ k) \quad [2]$$

$$Y_{j,t} \sim \begin{cases} MVN(\mu_t, \Omega_t) & Y_{L,j} \leq Y_j \leq Y_{U,j} \\ Y_{L,j}\ or\ Y_{U,j} & Y_{L,j} \geq Y_j or\ Y_j \geq Y_{U,j} \end{cases} \quad [3]$$

Where on day t, μ is the vector of estimated means, **Ω** is the estimated precision matrix, Y is the data from NMME, weather variable $j$, where any ensemble member that was below the lower bound ($Y_{L,j}$) or above the upper bound ($Y_{U,j}$) was set to the boundary. The priors for $\mu_t$ and $\Omega_t$ were set as follows: $\mu_0$ is a vector of zeros, **Σ** and $\boldsymbol{\Lambda}$ were set to a j x j diagonal matrix with 0.01, and k was set to j+1, where j is the number of weather variables. $Y_{L,j}$ and $Y_{U,j}$ are theoretical minimum and maximum values for each weather variable.

Cumulative growing degree days were calculated for each temperature ensemble member with a base of 10°C. We then used the mean (μ) and precision ($\tau$) across ensemble members to estimate daily cumulative growing degree days with error:

$$cgdd_t \sim N(\mu_t,\ \tau_t) \quad [5]$$

The hindcasts that use NMME have a horizon that is slightly less than 365 days because NMME, which has a one-year horizon, is produced once a month. For example, a hindcast starting on April 15th will use the NMME forecast starting on April 1st of that year, meaning the end of the hindcast is March 31st of the following year, giving a horizon of 350 days.

#### 1.2 Parameter updates

For all parameters in the survival and transition models (intercepts and coefficients tied to the effect of weather and mice), we assumed a Normal distribution and calculated the mean and precision of these parameters from the ensemble samples from the previous forecast, which became the hyperparameters for their prior distributions. For process error, we used moment matching by using the mean (μ) and variance (σ) from the posterior distribution of process error to estimate shape (α) and rate (β) hyperparameters for the inverse gamma priors on process error for the new forecast (Llera & Beckmann, 2016).

$$\alpha = \frac{\mu^2}{\sigma} + 2 \quad (6)$$

$$\beta = \mu * \left(\frac{\mu^2}{\sigma} + 1\right) \quad (7)$$

All parameters except reproduction (i.e. survival, transition, weather and mouse effect coefficients, process error), were updated at each iteration. Reproduction, which was never updated with empirical data, was held constant at the posterior mean from the calibration period to isolate the impact of larval data on forecast skill. This prevented the reproduction parameter from adjusting when larval data was omitted, ensuring a clear distinction between forecast performance with and without that specific data input.

### 2 Supplementary Results

#### 2.1 Larval forecasts

##### Skill vs lead time

**Figure S2.1**

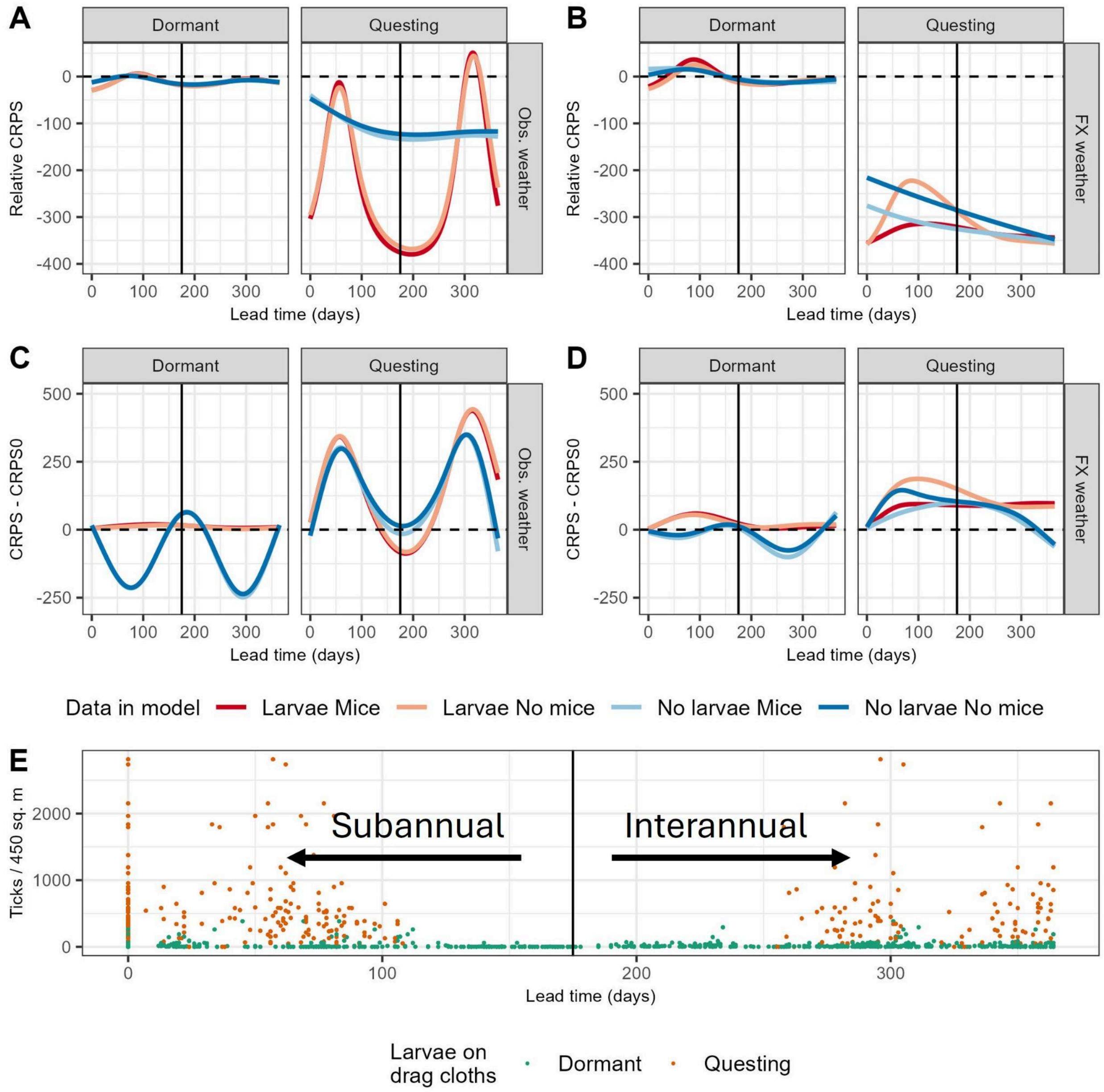

**Figure S2.1 –** Generalized additive models (GAMs) predicting the continuous ranked probability score (CRPS) of larval forecasts as a function of forecast lead time, split across forecasts for dormant and questing periods. A & B) Show GAMs predicting forecast CRPS, a measure of absolute skill, where A) depicts forecasts that used observed weather data (Obs. weather), and B) forecasts that used the forecasted weather data (FX weather). Values are relative to day-of-year forecasts, meaning values above zero indicate that the process models performed worse than the day-of-year model and values below zero indicate when the process models were more skillful. C & D) GAMs predicting the change in CRPS of larvae relative to the "nowcast" (day 0), a measure of predictability with respect to forecast lead time, where values above zero indicate that forecasts became less predictable with lead time, below zero they got more predictable. C) Forecasts that used observed weather data, and D) forecasts that used the forecasted weather data. E) Observed larvae on drag cloths as a function of lead time where color indicates if the tick drag occurred during the dormant (green) or questing (orange) period of the year. The vertical lines on day 175 represents the shortest lead time for interannual forecasts (the longest lead time for subannual forecasts was 178 days).

#### Forecast limit

**Table S2.1**

| Period | Larval data | Mouse data | Forecasted weather | Observed weather |
|---|---|---|---|---|
| Questing | Included | Included | > 365 | 303 |
| | | Excluded | > 365 | 304 |
| | Excluded | Included | > 365 | > 365 |
| | | Excluded | > 365 | > 365 |
| Dormant | Included | Included | 35 | 66 |
| | | Excluded | 46 | 67 |
| | Excluded | Included | 0 | 58 |
| | | Excluded | 0 | 56 |

**Table S2.1 -** Forecast limit (lead time in days) for larval ticks by forecast time scale, field data included, and weather data used in the model. The forecast limit reported is the mean forecast lead time in which the process-models became less skillful than the day-of-year model.

Transferability

**Figure S2.2**

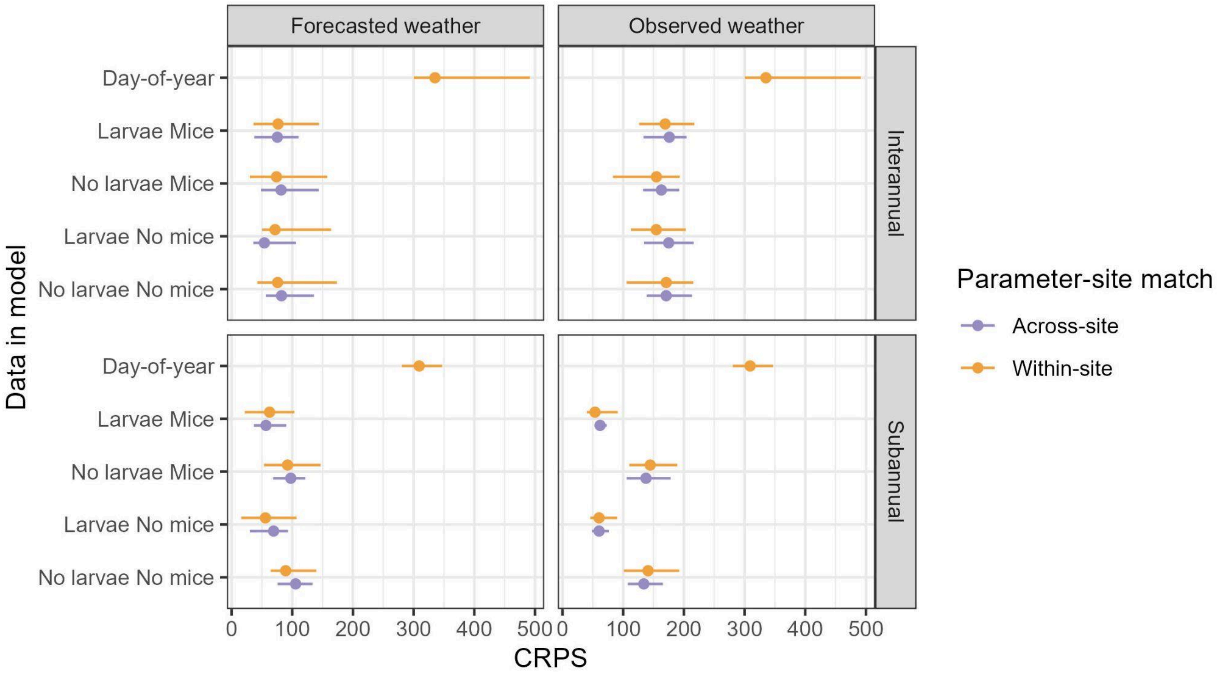


**Figure S2.2 -** CRPS 90% confidence intervals and medians (x-axis) for questing larvae given the data used in the model (y-axis), where forecasts that used parameters calibrated in the same site that was forecasted (within-site) are compared against forecasts that used parameters originally calibrated at one site and used to forecast ticks in another (across-site). The null day-of-year model was only run within-site.

#### 2.2 Adult forecasts

##### Skill vs lead time

**Figure S2.3**

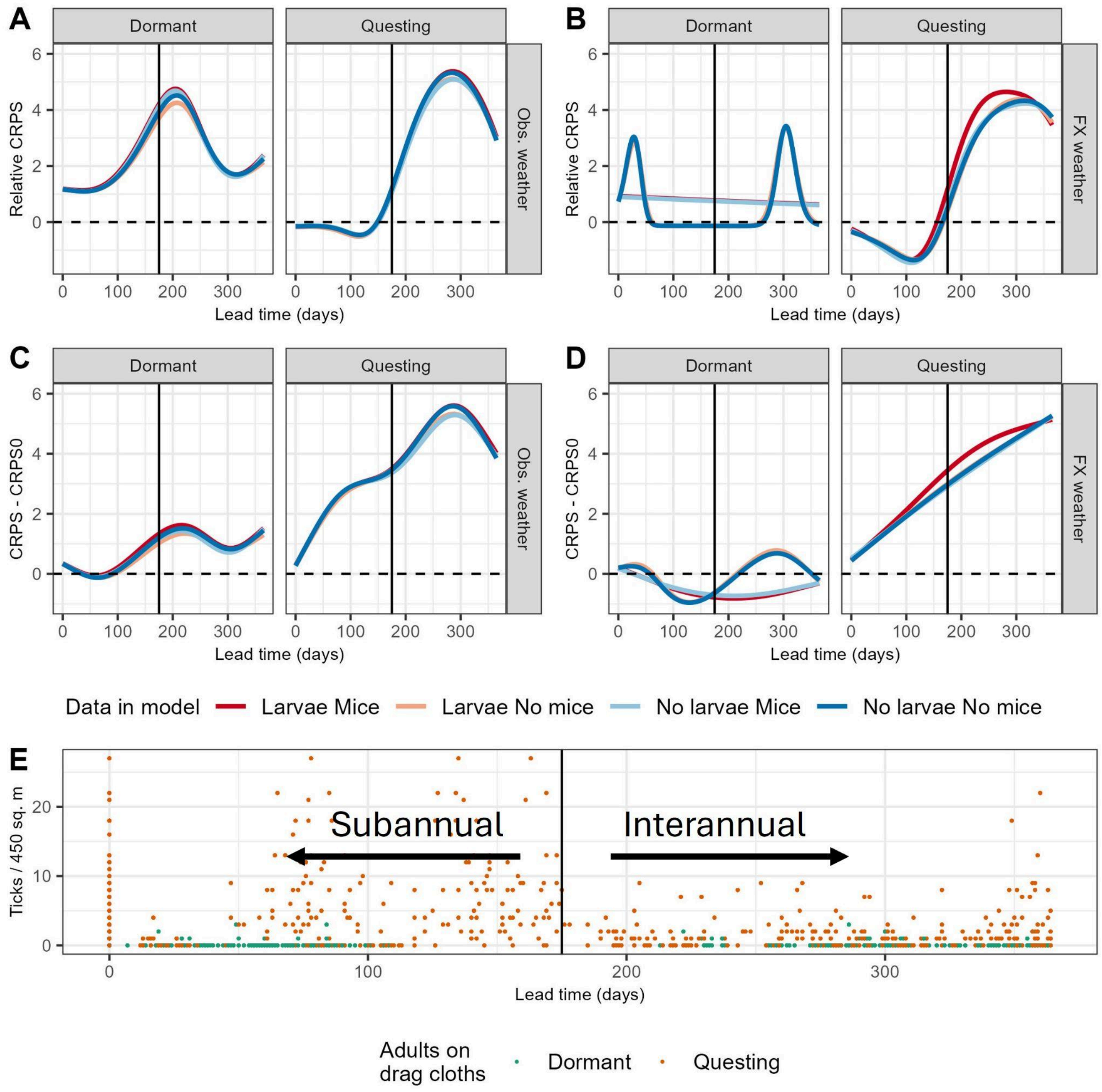


**Figure S2.3 –** Generalized additive models (GAMs) predicting the continuous ranked probability score (CRPS) of adult forecasts as a function of forecast lead time, split across forecasts for dormant and questing periods. A & B) Show GAMs predicting forecast CRPS, a measure of absolute skill, where A) depicts forecasts that used observed weather data (Obs. weather), and B) forecasts that used the forecasted weather data (FX weather). Values are relative to day-of-year forecasts, meaning values above zero indicate that the process models performed worse than the day-of-year model and values below zero indicate when the process models were more skillful. C & D) GAMs predicting the change in CRPS of adults relative to the “nowcast” (day 0), a measure of predictability with respect to forecast lead time, where values above zero indicate that forecasts became less predictable with lead time, below zero they got more predictable. C) Forecasts that used observed weather data, and D) forecasts that used the forecasted weather data. E) Observed adults on drag cloths as a function of lead time where color indicates if the tick drag occurred during the dormant (green) or questing (orange) period of the year. The vertical lines on day 175 represents the shortest lead time for interannual forecasts (the longest lead time for subannual forecasts was 178 days).

Forecast limit

**Table S2.2**

| Period | Larval data | Mouse data | Forecasted weather | Observed weather |
|---|---|---|---|---|
| Questing | Included | Included | 157 | 150 |
| | | Excluded | 168 | 150 |
| | Excluded | Included | 164 | 151 |
| | | Excluded | 167 | 149 |
| Dormant | Included | Included | 0 | 0 |
| | | Excluded | 0 | 0 |
| | Excluded | Included | 0 | 0 |
| | | Excluded | 0 | 0 |

**Table S2.2** - Forecast limit (lead time in days) for adult ticks by forecast time scale, field data included, and weather data used in the model. The forecast limit reported is the mean forecast lead time in which the process-models became less skillful than the day-of-year model.

Transferability

**Figure S2.4**

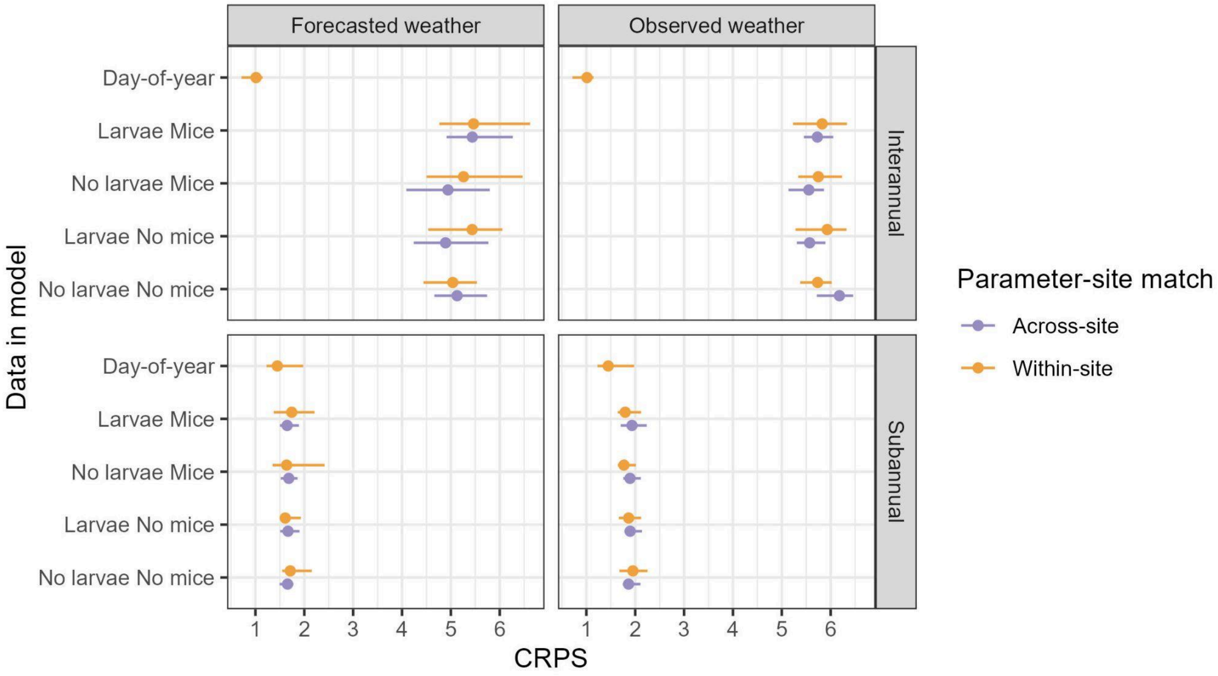


**Figure S2.4 -** CRPS 90% confidence intervals and medians (x-axis) for questing adults given the data used in the model (y-axis), where forecasts that used parameters calibrated in the same site that was forecasted (within-site) are compared against forecasts that used parameters originally calibrated at one site and used to forecast ticks in another (across-site). The null day-of-year model was only run within-site.

#### 2.3 Data assimilation

The data assimilation scheme updated states and parameters as expected. For example, one of the more skillful forecasts was issued on November 11, 2016 for June 22, 2017, and made a median forecast of 8.64 (90% CI 4.86-12.63) nymphs/450$m^2$ when the observed density was 9 nymphs/450$m^2$ (90% CI 4-14 assuming Poisson error). After the analysis step, the new initial condition for the next forecast had a median of 9.05 (90% CI 5.77-12.68) nymphs/450$m^2$ (Figure S3.3). Additionally, the variance for these distributions was 5.23, 9, and 4.43 for the forecast, data, and initial condition, respectively. Meaning that the forecast was more precise than the data, and, after the analysis step, the initial condition was more precise than the forecast.

By contrast, when a bad forecast was made, the analysis step shifted the initial condition distribution towards the data. For example, the forecast that was issued on August 1, 2012 for June 4, 2013 made a median prediction of 24.48 (90% CI 12.45-41.37) nymphs/450$m^2$ when the observed density was 58 nymphs/450$m^2$ (90% CI 46-71 assuming Poisson error). After the analysis step, the new initial condition for the next forecast had a median of 46.18 (90% CI 38.58-54.69) nymphs/450$m^2$ (Figure S3.3).

### 3 Supplementary Figures

**Figure S3.1 estimated latent weather**

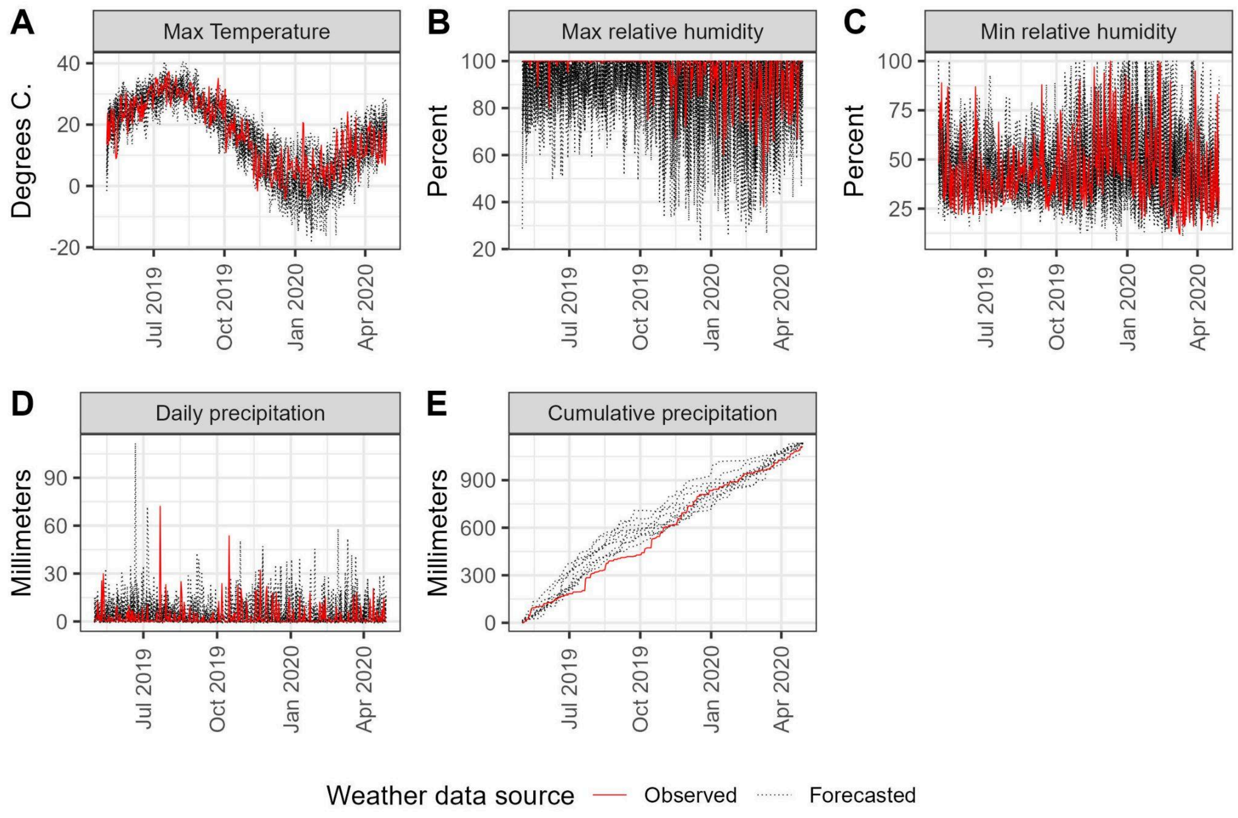


**Figure S3.1 -** An example of the NMME forecast vs observed meteorology at Cary. Each individual bias-corrected NMME forecast is in black, while the observed values from Cary are in red. The forecast was issued on May 20, 2019. Cumulative precipitation was calculated using the daily precipitation values for each ensemble.

**Figure S3.2**

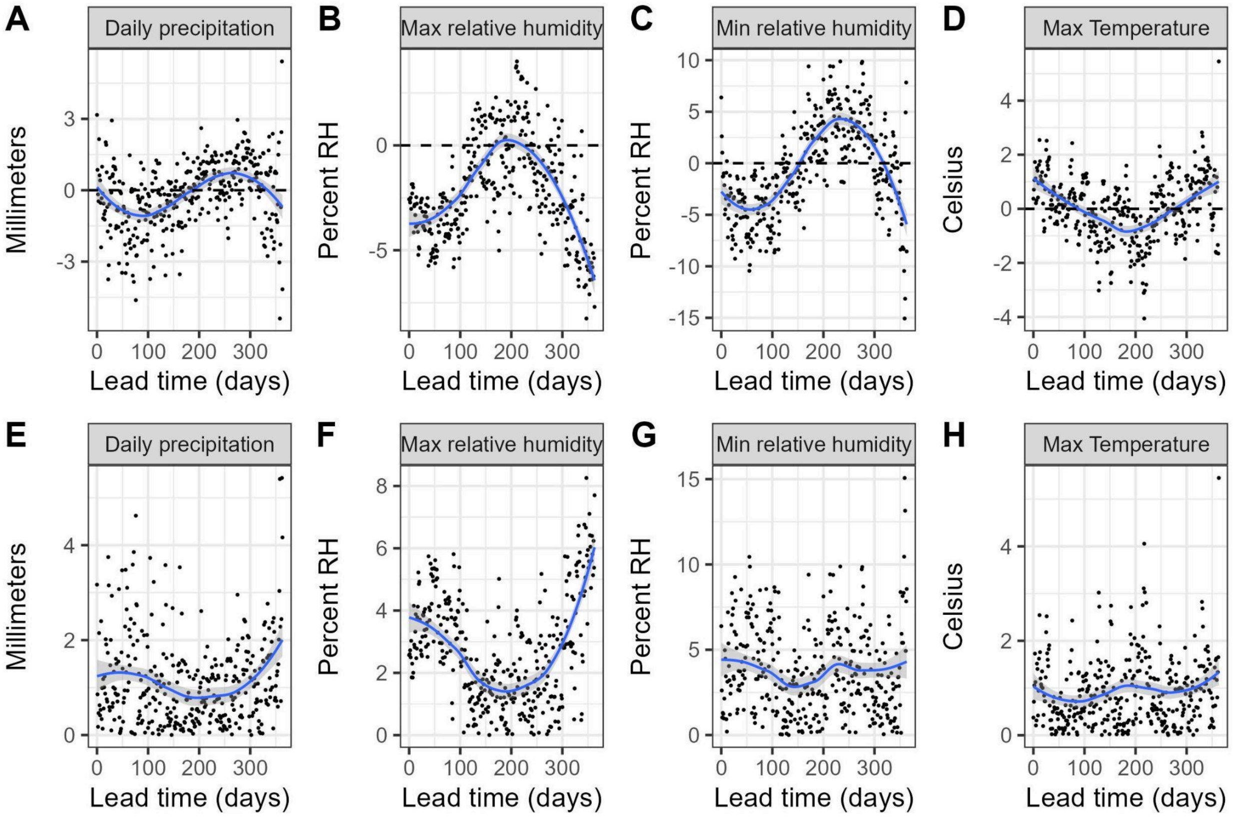


**Figure S3.2** - Top row (A-D) is the mean bias (forecasted weather minus observed weather) across forecast lead time in days for each weather variable used in the process models. Bottom row (E-H) is the mean absolute error of the forecast's weather compared to the observed weather. Loess curves (blue) to show trends. Bias and MAE were calculated after bias-correction of each ensemble member (see supplementary methods).

**Figure S3.3 Data assimilation**

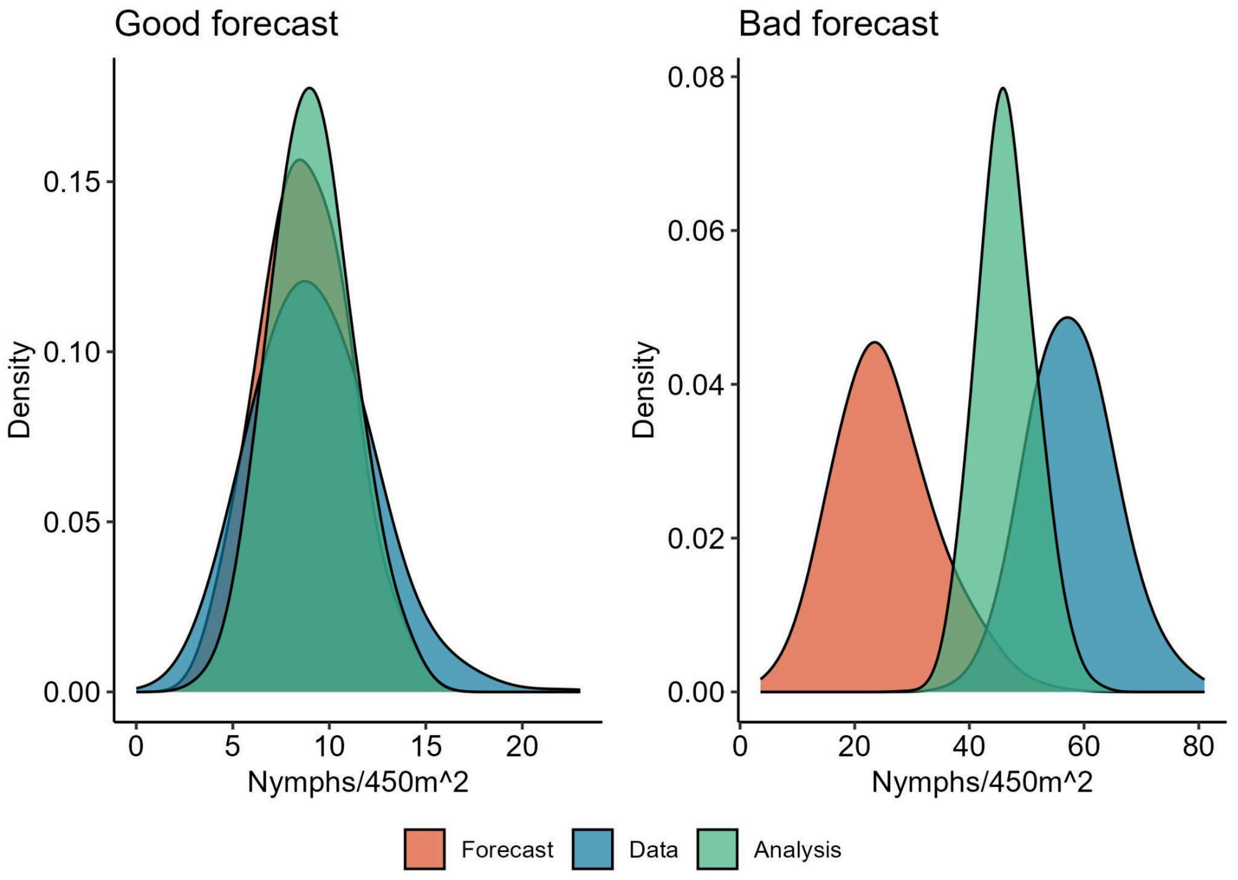


**Figure S3.3 -** How the data assimilation works for two nymph forecasts, a good forecast (issued on November 11, 2016 for June 22, 2017) on the left and a bad forecast (issued on August 1, 2012 for June 4, 2013) on the right. In order of occurrence, we first make a forecast (orange distribution). Then, we make an observation (blue), which is a Poisson distribution with a mean equal to the observed number of nymphs. Lastly, the analysis distribution (green) is the estimate of the latent state on the observation day given both the forecast and data, and becomes the initial condition for the next forecast.
